# Cortical depth-dependent human fMRI of resting-state networks using EPIK

**DOI:** 10.1101/2020.12.07.414144

**Authors:** Patricia Pais-Roldán, Seong Dae Yun, Nicola Palomero-Gallagher, N. Jon Shah

**Author notes:** **Correspondence:** Prof. N. Jon Shah. **Author Contributions** *N.J.S. designed the research; S.Y. developed the MR imaging sequence and the corresponding image reconstruction software; S.Y. and P.P. performed in vivo experiments. P.P. designed and performed data analysis; P.P. wrote the manuscript and prepared figures; N.J.S. invented the original EPIK sequence; N.P.-G. verified the anatomical labels and wrote the cytoarchitecture-related discussions; S.Y., N.P.-G. and N.J.S. reviewed the manuscript. **Data Availability Statement** *The original* in vivo *data from 13 subjects can be shared by submitting a formal request to the corresponding author (N. Jon Shah). The general protection of personal data and privacy policy, declared by the Council of Europe (https://www.coe.int/en/web/portal/personal-data-protection-and-privacy), applies to health-related data (CM/Rec(2019)2). The protection of* in vivo *data or metadata derived from the original data is further described by the corresponding ethics/internal administrating documents. The sharing is based on the consent of the subject whose data are to be shared; these subjects will be informed beforehand.

## Abstract

Recent laminar-fMRI studies have substantially improved understanding of the *evoked* cortical responses in multiple sub-systems; in contrast, the laminar component of *resting-state networks* spread over the whole brain has been less studied due to technical limitations. Animal research strongly suggests that the supragranular layers of the cortex play a critical role in maintaining communication within the default mode network (DMN); however, whether this is true in this and other human cortical networks remains unclear. Here, we used EPIK, which offers unprecedented coverage at sub-millimeter resolution, to investigate cortical broad resting-state dynamics with depth specificity in healthy volunteers. Our results suggest that human DMN connectivity is primarily supported by intermediate and superficial layers of the cortex, and furthermore, the preferred cortical depth used for communication can vary from one network to another. This has major implications for the characterization of network-related diseases and should occupy future high-resolution studies. In addition, the laminar connectivity profile of some networks showed a tendency to change upon engagement in a motor task. In line with these connectivity changes, we observed that the amplitude of the low-frequency-fluctuations (ALFF), as well as the regional homogeneity (ReHo), exhibited a different laminar slope when subjects were either performing a task or were in a resting state (less variation among laminae, i.e., lower slope, during task performance compared to rest). The varied laminar profiles concerning network connectivity, ALFF, and ReHo, observed across two brain states (task vs. rest) suggest the potential diagnostic value of laminar fMRI in psychiatric diseases, e.g., to differentiate the cortical dynamics associated with disease stages linked, or not linked, to behavioral changes. The evaluation of laminar-fMRI across the brain encompasses computational challenges; nonetheless, it enables the investigation of a new dimension of the human neocortex, which may be key to understanding neurological disorders from a novel perspective.

## 1. INTRODUCTION

In the mammalian brain, the cerebral cortex is organized into specialized functional areas across its surface. In humans, the largest part of the cerebral cortex is occupied by the neocortex, where the perikarya of multiple neuronal cell types are organized into six horizontal layers. The layered structure of the neocortex, which can be characterized by its cytoarchitecture, is involved in specialized signal processing (Palomero-Gallagher and Zilles, 2019, Thomson and Bannister, 2003, Larkum et al., 2018). The functional specificity of the cortex at its different depths has been demonstrated in animals using multi-site electrodes and, more recently, with cell-specific calcium imaging, e.g., via two-photon microscopy or optical fibers (Michelson and Kozai, 2018, Krupa et al., 2004, Adesnik and Naka, 2018, Ayaz et al., 2019, Scott et al., 2018). Although electrophysiology can theoretically be performed in the human brain to investigate layer-specific activations, it is an extremely invasive technique, and, consequently, data are only available from select patient groups, e.g., in epileptic patients, who benefit from such an undertaking (Csercsa et al., 2010). Moreover, although electrode recordings can provide direct information concerning layer activity, the coverage of electrophysiological recordings is limited by the number of multi-site electrodes, which is usually a single point at different depths. In contrast to the limitations of electrophysiology, functional magnetic resonance imaging (fMRI) can afford broad brain coverage non-invasively, although it is usually limited by relatively low temporal and spatial resolution. In rodents, experiments merging fMRI and axonal tracing maps demonstrated that layers 2/3, but not other cortical depths, project exclusively to areas within the default mode network (DMN) (Whitesell et al., 2021), and it has been shown that stimulation of upper, but not lower, sections of the motor cortex results in network-wide effects (Weiler et al., 2008), suggesting a primary role of the superficial cortical laminae in resting-state networking. Due to a lack of non-invasive methods to sufficiently sample the cerebral cortex, the specific way in which the cortical neurons communicate with each other in the human brain -across different depths and through different areas-largely remains an open challenge, which could be addressed with novel high-resolution fMRI methods.

fMRI most commonly relies on the identification of blood changes, which, assuming perfect neuro-vascular coupling, represent neuronal function in the brain (Logothetis et al., 2001, Logothetis, 2002, Ogawa et al., 1990). Despite its dependence on the vascular architecture of the cortex, laminar fMRI, i.e. sub-millimeter resolution fMRI acquired with the purpose of studying depth-specific neuronal responses, has proven to be a valuable tool for studying cortical dynamics in animals and humans (for a review of methods and applications, see references (Lawrence et al., 2019, Petridou and Siero, 2019, Scheeringa and Fries, 2019, Huber et al., 2021a)). Two main fMRI contrasts are usually employed in human laminar fMRI: cerebral blood volume (CBV), typically applied as part of vascular occupancy schemes (VASO), and blood oxygenation level dependent (BOLD). Although VASO offers high spatial specificity by detecting dilation of the microvasculature based on T1 contrast, it exhibits low sensitivity due to unwanted tissue suppression by an inversion recovery pulse (Lu et al., 2003, Huber et al., 2018, Poplawsky et al., 2015). In contrast, sequences based on BOLD present the advantage of a higher signal-to-noise ratio (SNR), with spin-echo-based (SE) echo planar imaging sequences (EPI) offering greater parenchymal specificity and gradient-echo (GE) schemes offering the best sensitivity [27]. The higher SNR and the flexibility of being able to accommodate higher sampling rates make GE sequences some of the most commonly used in the fMRI field, typically in the form of GE-EPI. Due to GE schemes having enhanced sensitivity to large vessels (Kay et al., 2019), with long T2* constants, e.g., pial veins, several methods have been presented for use in the pre-processing pipeline to ameliorate contamination from extra-parenchymal signals or to correct the evoked activation profile across cortical depths (Menon, 2002, Curtis et al., 2014, Fracasso et al., 2018, Kashyap et al., 2018a, Heinzle et al., 2016, Markuerkiaga et al., 2016, Kay et al., 2020).

fMRI acquired with different contrasts and sub-millimeter resolution has enabled the function of the cerebral cortex to be sampled with column- and layer-specificity in the context of sensory, motor, visual, and auditory tasks (Guidi et al., 2016, Huber et al., 2017, Chai et al., 2020, Huber et al., 2015, Siero et al., 2011, Yu et al., 2014, Ress et al., 2007, Polimeni et al., 2010, Kashyap et al., 2018b, van Mourik et al., 2019, De Martino et al., 2015, Muckli et al., 2015, Yacoub et al., 2008) and has shown the superficial layers of the neocortex to be predominantly involved in the processing of neuronal signals. This predominance is possibly due to the fact that layer III is the main source and target of cortico-cortical connectivity (Zilles and Catani, 2020) and that the higher synaptic density found there compared to the remaining cortical layers is associated with higher receptor densities (Palomero-Gallagher and Zilles, 2019). Importantly, a study focusing on the visual cortex reported correlations between the function of specific layers imaged with fMRI and particular EEG rhythms (Scheeringa et al., 2016), reinforcing the potential for using fMRI to detect layer variability.

In contrast to task-related circuits, which are usually limited to specialized responses in a small portion of the cerebral cortex, during rest, cross-talk between distant cortical areas maintains a baseline neurological state (Raichle et al., 2001). Although the potential for using fMRI to sample cortical layers is clear, the limited brain coverage enforced by most existing high-resolution-fMRI schemes precludes the investigation of large-scale inter-regional depth-dependent connections; i.e. some resting-state evaluations have been presented previously, but these mostly focused on particular systems, e.g., (Polimeni, 2010, Huber et al., 2017). To date, only a few groups have reported fMRI acquisitions from the whole brain at ≤ 1 mm resolution (De Martino et al., 2011, Sharoh et al., 2019, Huber et al., 2020), and, to our knowledge, only one study has intended to use laminar fMRI to uncover the cross-layer functional interactions that exist in the human brain during rest with a nominal spatial resolution of 0.8 × 0.8 × 0.8 mm^3^ (Huber et al., 2020).

Here, we applied TR-external EPI with keyhole (EPIK) (Yun et al., 2022) to identify the laminar signals from most of the cerebral cortex with a 0.63 mm iso-voxel size in healthy volunteers. It has been previously shown that EPIK (Shah, 2003, Shah NJ, 2004, Zaitsev et al., 2001, Zaitsev et al., 2005, Yun, 2020, Yun and Shah, 2020, Yun et al., 2022) can offer higher spatial resolution in dynamic MR studies with comparable hemodynamic-response detection compared to routine EPI (Yun et al., 2013, Yun and Shah, 2017, Caldeira et al., 2019, Shah et al., 2019, Yun et al., 2019). Moreover, the combination of TR-external phase correction with EPIK has been shown to further enhance spatial resolution (Yun, 2022). In this work, TR-external EPIK covered the human cerebrum almost entirely, with 123 slices sampling voxel volumes of ~ 0.25 mm^3^. The combination of a smaller voxel size and near whole-brain coverage is a key feature of this work when compared to previous laminar fMRI studies. By adding pre-processing steps to ameliorate the potential bias of the GE sequence, the acquired images allowed us to perform cortical depth-dependent analysis of broad resting-state networking in the human brain.

## 2. MATERIALS AND METHODS

### 2.1. Subjects

The functional data from thirteen healthy volunteers (eleven males and two females; age, 23-47 years; 12/13 right-handed, 1/13 left-handed (Schweisfurth et al., 2018)) were included in this study. Although eighteen subjects were originally measured, four of them were removed from the study due to the lack of/non-proper physiological recordings, and an additional set of data was removed after detecting high levels of motion. Hence, the structural scans of eighteen subjects were used for the anatomical study of the cortex, but only data from thirteen subjects were included in the functional analysis relevant to the majority of the results shown in this manuscript. Subjects were scanned in one single session for a period of ~55 min, which included two resting-state fMRI scans (‘rs-fMRI-1’ and ‘rs-fMRI-2’), one finger motor task fMRI scan (‘task-fMRI’), and an MP2RAGE scan. Due to poor anatomical-functional co-registration, the rs-fMRI-2 data from one subject were omitted, and another subject was excluded from both the rs-fMRI-2 analysis and the whole-brain laminar evaluations of the task-fMRI for the same reason (i.e. the number of subjects used in the whole-brain analysis were 13, 11 and 12 for rs-fMRI-1, rs-fMRI-2, and task-fMRI, respectively). However, the task-fMRI data from all 13 subjects were used for the analysis of evoked responses in the primary motor cortex (M1) as the motor cortex could be well co-registered in all subjects. A pneumatic belt was positioned around the subject’s chest, and a pulse oximeter was placed on the 2^nd^, 3^rd^, or 4^th^ finger of the left hand to control for potential interference of the physiological signals on the fMRI responses (Glover et al., 2000). The experimental methods were approved by the local institutional review board (EK 346/17, RWTH Aachen University, Germany), MR-safety screening was performed prior to the MRI acquisition, and informed written consent was obtained from all subjects.

### 2.2. Experimental design

In the rs-fMRI, subjects were asked to remain still, awake, with eyes closed, and without thinking about anything in particular. In total, 172 volumes were reconstructed from the rs-fMRI (~10 min). For task-fMRI, subjects were asked to follow instructions on a projected screen and to perform flexion of the right index finger without necessarily touching the thumb. The task was performed continuously at a comfortable pace (1-2 Hz, for most subjects) for 21 s and was repeated 12 times, alternated with 21 s of rest. In total, 148 volumes were obtained (4 s rest, [21 s task, 21 s rest] × 12) (~ 8.6 min). One subject was invited to participate in an additional scan session and first performed a finger motor task (4 s rest, [21 s task, 21 s rest] × 8), followed by a finger motor and sensory task, i.e. instructions were given to approach the right index finger towards the thumb involving touching (4 s rest, [21 s task, 21 s rest] × 8).

### 2.3. MRI data acquisition

MRI data were collected on a Siemens Magnetom Terra 7T scanner with a 1-channel Tx / 32-channel Rx Nova Medical head coil. An anatomical MRI volume was acquired using an MP2RAGE sequence, TR/TE = 4300/2 ms, matrix = 256 × 376 × 400 (0.60 × 0.60 × 0.60 mm^3^). Functional MRI data were obtained using GE-EPIK combined with a TR-external EPI phase correction scheme. The sequence performance has been previously described elsewhere and briefly consists of a full sampling of central k-space (48 central lines) and a sparse sampling of the peripheral k-space (288 lines), which is completely acquired after three TRs, i.e. every TR encompasses the acquisition of the k-space center and 1/3 of the periphery (details of the sequence can be found in previous publications (Yun and Shah, 2017, Yun et al., 2013, Yun et al., 2019, Yun and Shah, 2020, Yun, 2022, Yun et al., 2022)). Sampling the central k-space at every TR ensures a near-optimum SNR for each individual scan. An analysis of raw data from a healthy subject in our preliminary study showed that the central k-space in the current configuration has 86.5% of the entire k-space energy, whereas including the peripheral k-space lines that are continuously updated at every scan (i.e. 1/3 of the periphery) results in 91.0% of the entire energy. Two acquisition protocols were used to collect data in this study. In both protocols, TR/TE = 3500/22 ms, FA = 85°, partial Fourier = 5/8, 3-fold in-plane/3-fold inter-plane (multi-band) acceleration, bandwidth = 875 Hz/Px, and αPC/αMain = 9°/90°. B0 shimming was performed with a standard routine provided by the manufacturer. For protocol-1 (used to generate all the data in the main figures), matrix = 336 × 336 × 123 slices (0.63 × 0.63 × 0.63 mm^3^). For protocol-2, matrix = 408 × 408 × 105 slices (0.51 × 0.51 × 1.00 mm^3^). The main figures show results acquired with protocol-1 (i.e. isotropic voxels).

### 2.4. Data analysis

A summarized pipeline of our pre-processing methods can be found in **Fig. S1**. Briefly, cortical surfaces were extracted from the anatomical scan following an equi-distance sampling approach and used as a template to map the functional signals using Freesurfer (Martinos Center for Biomedical Imaging, Charlestown, MA). An equi-volume layering sampling was additionally employed (LAYNII, https://github.com/layerfMRI/LAYNII, (Huber et al., 2021b)) to compare the functional results relevant to gyri/sulci (the equi-distance model applies to all figures shown in this report unless specified otherwise). The assessment of the cortical model (results in **Fig. S2**) is based on the work by Kay et al. (Kay et al., 2019). Pre-processing included the correction of the magnitude time course to minimize large-vein contribution (**Fig. S3**), as previously described (Menon, 2002, Curtis et al., 2014, Stanley et al., 2021), in conjunction with common pre-processing steps (slice-timing correction, realignment, temporal filtering, and regression of motion parameters as well as averaged white matter and CSF time courses) (Pais-Roldán, 2022), which resulted in zero-mean pre-processed functional time courses. For the assessment of the resting-state measures in task-related scans, the task predictor time course was added as a regressor-of-no-interest so as to avoid interference from an active motor pathway on the assessed metrics (for comparison, the power spectrum and connectivity measures were assessed in all conditions, including the task scan with and without regression of the task paradigm). Post-processing analysis included volume-based ICA-analysis, surface-based temporal correlation (i.e. functional connectivity), surface-based spectral decomposition, amplitude of low-frequency fluctuations (ALFF), surface- and volume-based regional homogeneity (ReHo), and, in task fMRI only, surface-based general linear model (GLM) analysis. Post-processing was performed using 3dRSFC and 3dReHo on AFNI (Analysis of Functional NeuroImages, NIH, Bethesda, MD), Matlab scripts (Mathworks, Natick, MA), and melodic on FSL (FMRIB Software Library, Oxford, UK). The implementation of surface ReHo was based on the functions of the Connectome Computation System ((Xu, 2015)). Volume-based analyses (ALFF, volume-based ReHo, and ICA) were followed by a volume-to-surface projection and averaging of vertices within each ROI for a total of 42 ROIs (21 per hemisphere, see **Table 1**) and each surface. Normalization of functional connectivity, ALFF, and ReHo was performed by subtracting the mean value across the brain and dividing by the standard deviation. Computations involving sulcal or gyral locations were restricted to vertices corresponding to the top or bottom 20% curvature values, respectively.

**Table 1.** Regions of interest. DMN: default mode network; EXEC: executive control network; SM: sensory-motor network. ROIs extracted from the Freesurfer surface parcellation *aparc.a2009s.annot*, based on the Destrieux atlas.

| Abb. ROI | Full ROI name | Related network(s) |
| --- | --- | --- |
| MPCC | Posterior Midcingulate Cortex | DMN |
| ACC | Anterior Cingulate Cortex | DMN |
| PCC | Posterior Cingulate Cortex | DMN |
| PCUN | Precuneus | DMN & VISUAL |
| STS | Superior Temporal Sulcus | DMN & EXEC |
| PoCG | Postcentral Gyrus | EXEC & SM |
| ANG | Inferior Parietal - angular | EXEC |
| SMG | Inferior Parietal - supramarginal | EXEC |
| IFG | Inferior Frontal G&S | EXEC |
| MFG | Middle Frontal Gyrus | EXEC |
| SFG | Superior Frontal G&S | EXEC & SM |
| SOG | Superior Occipital Gyrus | VISUAL |
| POS | Parieto-occipital Sulcus | VISUAL |
| CAL | Calcarine | VISUAL |
| CUN | Cuneus | VISUAL |
| CS | Central Sulcus | SM |
| PreCG | Precentral Gyrus | SM |
| PreCS | Precentral Sulcus | SM |
| STG | Superior Temporal Gyrus | AUDITORY |
| SubCG | Subcentral G&S | AUDITORY |
| MACG | Anterior Midcingulate Cortex | SALIENCE |

Note that the term “surfaces” or “layers” employed through the manuscript should not be interpreted as histologically-defined cortical layers; instead, the terms are employed to refer to each of the six different cortical depths that are artificially generated by splitting the cortical thickness into five different compartments. A more detailed explanation of the methods used to generate the presented results can be found in the *Supplementary Material*.

## 3. RESULTS

### 3.1. Characterization of the cortical template for whole-brain laminar mapping

In a recent report, we demonstrated the feasibility of using TR-external EPIK to assess fMRI activity over a large FOV with submillimeter spatial resolution (Yun, 2020). The additional acquisition of a structural MP2RAGE sequence enabled us to produce a surface model of the cerebral cortex for each subject. Based on this model, it was possible to study brain activity at six different cortical depths, referred to hereafter as “layers” (different from the cytoarchitectonically-defined cortical layers) and specified as “pial, +80%, +60%, +40% +20%, and white”, from the most superficial to the deepest portion of the cortex, respectively. These layers were used as a template for analysis of the functional data (see *Methods* and **Fig. S1**). **Fig. S2** provides a characterization of the cortical model. In **Fig. S2A**, an example of the distribution of sulci and gyri, i.e. the curvature of the cerebral cortex and cortical thickness, is provided for a representative coronal slice. The spatial resolution of each surface was assessed in sulci and gyri independently. As the cortical surfaces were generated by inflating the inner cortical boundary towards the pial space, a one-to-one correspondence between vertices existed across all surfaces; hence, the in-surface resolution varied depending on the depth and the curvature (for a visual example, see insets in **Fig. S2B**). The histograms in **Fig. S2B** show the distribution of vertex-to-vertex distances at different cortical depths for gyri (blue) and sulci (green), in both cerebral hemispheres, as a mean ± standard deviation (SD), obtained from an 18-subject group. In agreement with previous work (Kay et al., 2019), it was observed that, in gyri, the vertex-to-vertex distance increased from ‘white’ to ‘pial’; on average, a sparse distance of 0.72±0.04 mm existed between the vertices in the pial surface, whereas this distance was 0.49±0.01 mm in the white surface, i.e. the surface spatial resolution improved with depth in gyri locations. In contrast, in sulci, the vertex-to-vertex distance decreased from ‘white’ to ‘pial’, with an average of 0.36±0.01 mm between consecutive vertices in the pial surface vs. 0.49±0.02 mm between the vertices of the deepest layer, i.e. outer surfaces were more densely sampled in sulci. This feature was observed in both the left and right hemispheres. The upper inset in **Fig. S2B** exemplifies the larger vertex-to-vertex distance in the superficial layers of the cortical gyri, and the inset below exemplifies the larger distance between consecutive vertices in gyri compared to sulci. Cortical thickness was calculated in both cerebral hemispheres for 21 ROIs and for gyri and sulci in the whole cortex (**Fig. S2C**). Note that the analysis specific to gyri or sulci only considered the bottom 20% and top 20% curvature values, respectively (see *Methods*). Given that the cortical ribbon is thicker in the crown of gyri than in their walls or in the fundus of sulci (Kay et al., 2019) (**Fig. S2C**, blue bar vs. green bar), the layer-to-layer separation is greater at this position of the gyri (**Fig. S2D**, 0.62±0.02 mm between layers of cortical gyri vs. 0.40±0.01 mm in sulcal locations), which indicates that higher signal contamination from neighboring layers can be expected in sulci, particularly in their fundus. Note that the in-depth resolution in the crown of gyri, i.e. the inter-layer distance, approximately corresponds to the voxel size employed in our fMRI protocol. The relative differences in surface resolution and inter-layer distance across cortical depths are in broad agreement with the study of Kay et al. (Kay et al., 2019). However, their results showed a slightly better surface resolution, which was probably due to having increased the density of surface vertices during processing, a step that was avoided here to limit the computational demands of the whole-cortex functional analysis. **Fig. S4** shows the sampling of the different cortical depths overlaid on the anatomical and the mean functional images for a representative case. The characterization of the cortical surfaces suggested that the chosen subject-specific cortical model could be used as a reliable template to map functional signals and to study layer-based functional oscillations.

### 3.2. De-vein of the high-resolution GE-EPIK

Given the susceptibility of GE images to the signal of ascending venules and pial veins, we adopted the correction method described by Menon et al. (Menon, 2002, Curtis et al., 2014). This method consists of using the information in the phase signal, which is presumably proportional to changes in oxygenated/deoxygenated hemoglobin in non-randomly oriented vessels, to remove unwanted signal change from the magnitude signal. The procedure should, therefore, minimize the contribution of large vessels (mainly in upper cortical layers and veins in the CSF compartment) while preserving the signal of the gray matter capillaries. **Fig. S3** exemplifies the correction method by displaying the average magnitude, phase, and corrected magnitude signal time course in four regions at different cortical depths in the precentral gyrus during task performance (the same method was applied to the resting-state data). The signal from voxels near the CSF (“1” in **Fig. S3C**) is substantially affected by Menon’s de-vein procedure, but the change is much less evident in deeper gray matter voxels (e.g., “3”, “4” or “5” in **Fig. S3C**). The contribution of large vessels can be reduced by using this correction step.

### 3.3. Depth-dependent evoked responses in the primary motor cortex (M1)

In order to prove the capabilities of TR-external EPIK for use in a laminar analysis of the whole brain, we first aimed to reproduce the results obtained in previous fMRI studies of finger-motor tasks. These results are characterized by a strong response in the upper layers of M1 and a second activation peak in the middle-deep layers when avoiding non-corrected GE-BOLD sequences (Huber et al., 2017, Chai et al., 2020, Huber et al., 2015). A task-based fMRI protocol consisting of 12 epochs, alternating 21 s blocks of finger motion with 21 s blocks of rest, was conducted on each subject using a submillimeter-resolution protocol configured with TR-external EPIK; voxel size: 0.63 mm isotropic, matrix: 336 × 336, 123 slices (see protocol-1 in *Methods*). In agreement with the previous literature (Huber et al., 2017, Chai et al., 2020, Huber et al., 2015), we observed a marked predominance of the BOLD response in the superficial layers of M1 (**Fig. 1**). A monotonous signal decrease from superficial to deep layers, typical in laminar GE-BOLD, was observed after averaging the beta coefficient within a portion of M1 (**Fig. 1A-B**, N=13), possibly reflecting a bias due to ascending veins and large venous vessels located on the surface (**Fig. S5**) that may remain even after the de-vein procedure. However, when sampling the beta-map of area M1-4A (Geyer et al., 1996) (specifically, along 20 lines, each extending from the CSF and up to the WM), we were able to identify a bimodal response superimposed over this gradient in multiple subjects and through several consecutive slices (see mean line profile in **Fig. 1C** and several examples through **Fig. S6**). This laminar activation profile resembled the typical profile of finger motor tasks finely assessed in high-resolution schemes (Huber et al., 2017, Chai et al., 2020, Huber et al., 2015). The specificity of laminar responses was further evaluated in one subject by comparing the activation profiles of two different tasks: movement-only, or movement + touch, i.e. somatomotion, or movement + somatosensation (**Fig. 1K**). While both tasks resulted in an initial activation peak in the superficial layers, the relatively salient response observed in deeper layers during movement was minimized and shifted towards more superficial layers upon the addition of touch. It is worth noting that the post-hoc correction steps to ameliorate the GE-BOLD signal bias towards pial vessels did not include deconvolution of model signal bias across the cortical ribbon (in order to keep consistent pre-processing across task and resting-state data). Although the pre-processed data are likely to be partially biased towards the surface, these results demonstrate the potential of TR-external EPIK to track laminar changes in the human cortex, even without the application of deconvolution procedures.

**Fig. 1.**
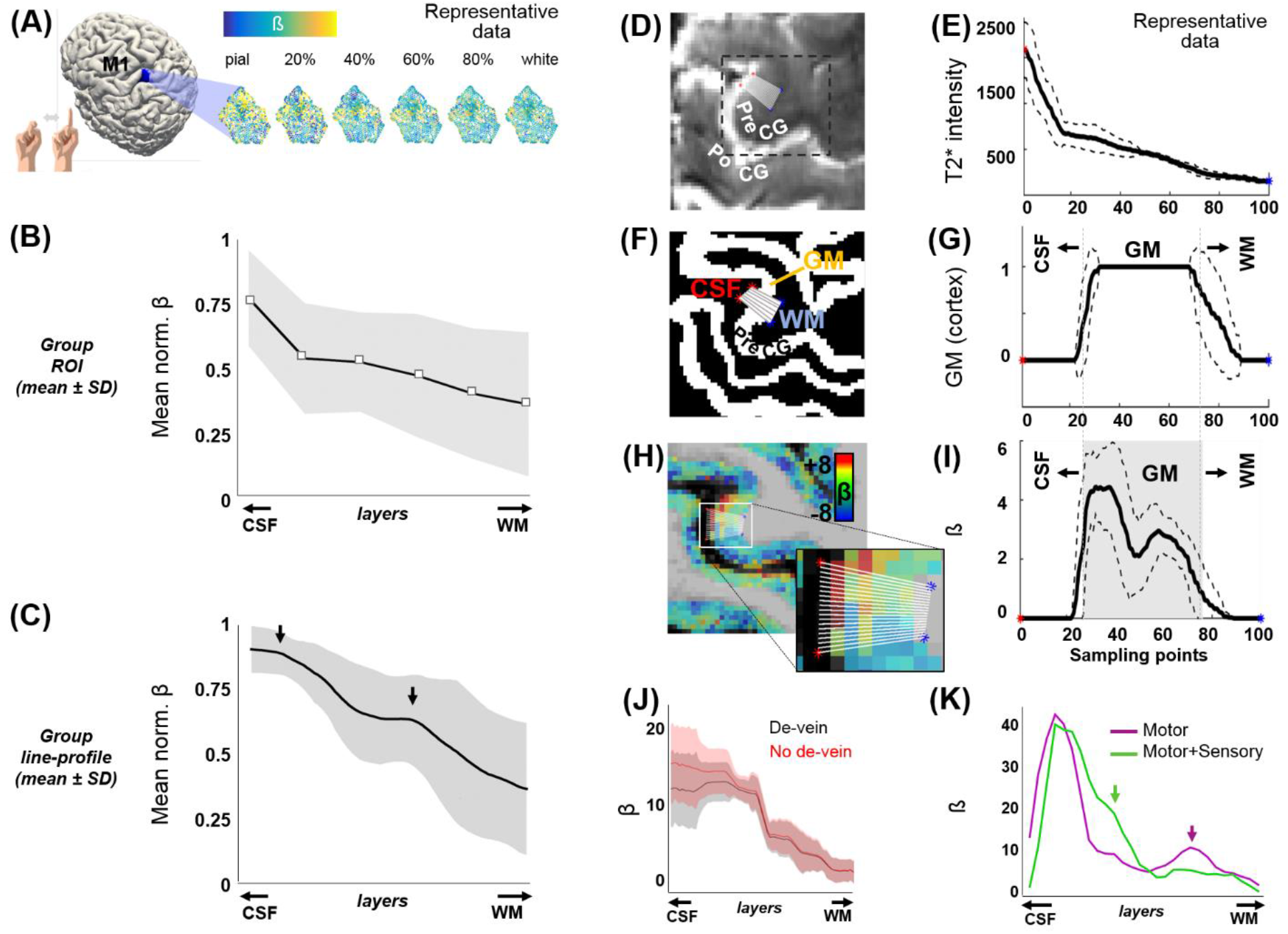
Cortical depth-dependent motor task-evoked BOLD fMRI. **(A)** Flattened patches of M1 extracted from six surfaces at different cortical depths in one volunteer. **(B)** The plot shows the mean ±SD of the normalized beta-coefficient in six cortical patches of M1 at different cortical depths (N=13). **(C)** Mean ±SD of the normalized activation line-profiles, computed from the beta map of 13 subjects, using a line-profile method on a particular segment of M1 (computation explained through panels (D)-(I)) The arrows point to the two main activation peaks observed along the cortical depth. **(D)** The figure shows part of a mean TR-external EPIK axial slice containing the precentral gyrus (PreCG). The 20 overlaid sampling lines (in white) extend from the CSF, covering the cortical thickness, up to the WM. **(E)** Average line profile of the mean TR-external EPIK on the selected sampling region. Note the high intensity at the initial points (CSF, with long T2 constant). The dashed lines above and below the solid line represent the mean ±SD. **(F)** Same region as in (D), cropped from the segmented MP2RAGE, which identifies the cortical ribbon (in white color, labeled as GM). **(G)** Line profile calculated from the sampling lines over the image in (F). **(H)** Beta coefficient map overlaid on the MP2RAGE and zoomed view of the 20 sampling lines. **(I)** Profile of the beta coefficient along the selected lines crossing the finger motor area in the PreCG. Note the higher signal on superficial and middle layers of the cerebral cortex (double peak, typical of motor tasks). The gray-shaded area in i) represents the extent of the GM, with superficial layers (near the CSF) on the left and deep layers (near the WM), on the right. **(J)** An example of an evoked line profile computed before (red) and after (black) the addition of the de-vein correction step. **(K)** Line profiles in a subject performing either finger movement only (purple trace) or finger movement involving touch (green trace). In panels (E), (G), (I), and (K), the x-axis refers to the distance from the CSF to the WM, equidistantly sampled with 100 points.

### 3.4. Near whole-brain assessment of cortical depth-dependent activity in resting-state fMRI

To investigate whole-brain depth-dependent functional measures and cross-layer interactions between distant cortical areas during rest, we acquired two additional task-negative fMRI scans from each subject, one using protocol-1, with 0.63 mm isotropic voxels, i.e. same as for the task-fMRI, and another using protocol-2, with higher in-plane resolution (0.51 × 0.51 mm^2^) and thicker slices (1.00 mm) to maintain a reasonable SNR level. In order to reduce the dimensionality of our data (6 surfaces × ~ 290 000 vertices per hemisphere), we first segmented the cerebral cortex into 21 ROIs related to well-known brain networks: default mode, visual, sensory-motor, auditory, executive control, and salience (**Table 1**). Having averaged the vertices within each ROI and each cortical layer in both cerebral hemispheres, we obtained 252 time courses from each fMRI scan (21 ROIs × six surfaces × two hemispheres). These were then subjected to several analysis methods. Following a frequency-power decomposition of the functional time courses, it was found that most cortical areas exhibit a higher level of activity in their superficial layers (**Fig. 2A**). This activity was observed to oscillate mostly within the range 0.01-0.03 Hz, which is in agreement with other non-laminar rs-fMRI studies focusing on cortical gray matter (Zuo et al., 2010, Xue et al., 2014, Yuen et al., 2019, Bajaj et al., 2014). Integration of the signals fluctuating between 0.01 and 0.1 Hz, i.e. the typical range considered in rs-fMRI studies (Murphy et al., 2013), provided a simplified measure of brain activity during rest, commonly denoted as ALFF, which was assessed for each ROI and the six different cortical depths. **Fig. S7** shows the averaged ALFF (across all ROIs and subjects) for each cortical depth, considering either the whole surface, only gyral locations, or only sulcal locations, and using either the data sampled with an equi-distance or an equi-volume sampling model. ALFF laminar profiles were generated for each ROI. To reduce dimensionality, a linear fitting of the laminar profile was computed, and the slope of the fitting line was calculated and reported per ROI (color-coded in **Fig. 2B**) and per network (**Fig. 2C**, see network color-coding in **Fig. 2D)**). The level of activity (ALFF) was observed to decrease from the surface of the cortex towards the deep gray matter in most ROIs (i.e. negative slope, hot colors in **Fig. 2B**), irrespective of whether these were assessed in cortical gyri or cortical sulci (**Fig. S7**). The sensory-motor network and the auditory network exhibited the most and the least steep ALFF laminar profiles, respectively, in both cerebral hemispheres (red and purple markers in **Fig. 2C**).

**Fig. 2.**
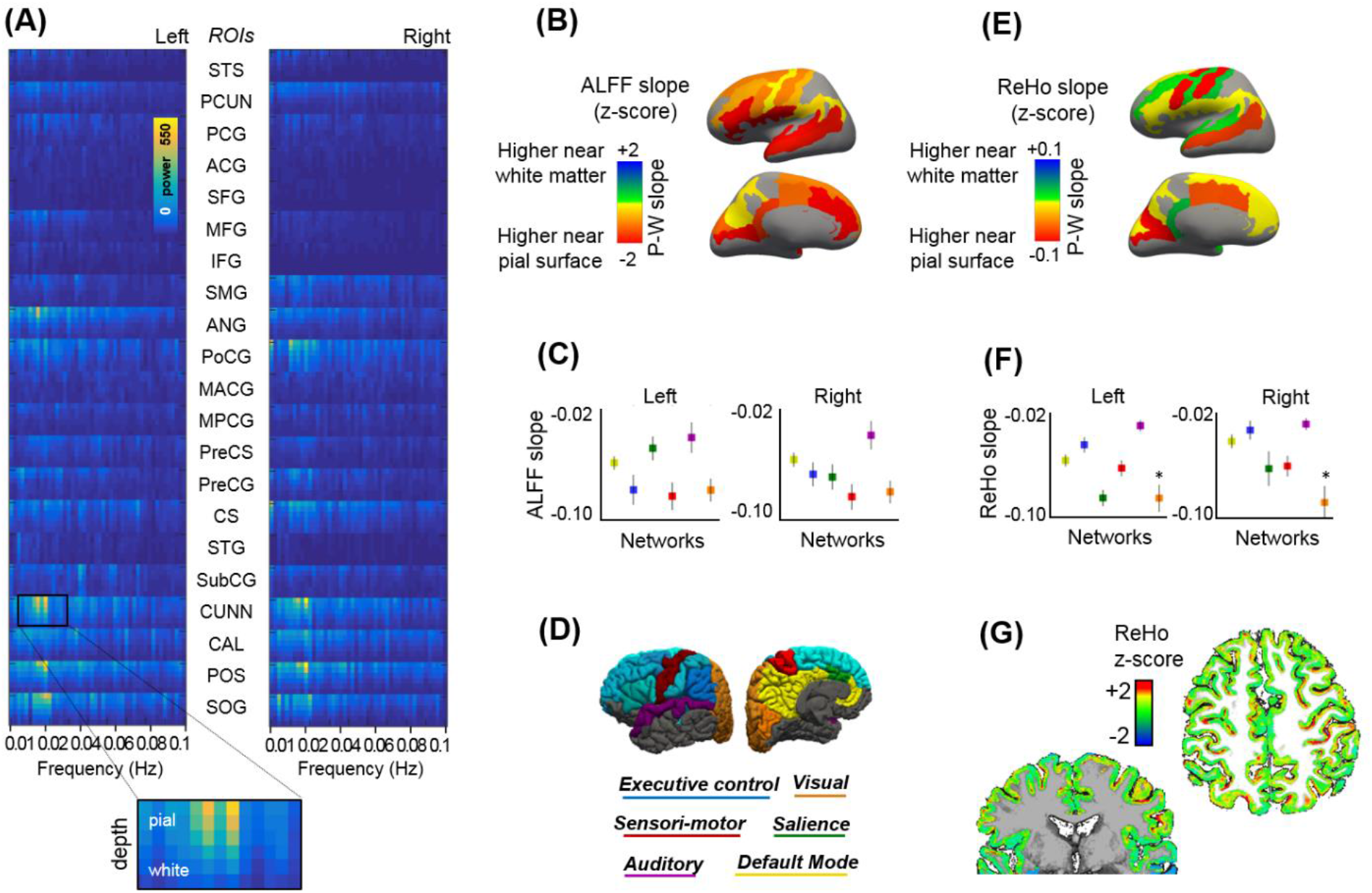
Cortical depth-dependent resting-state activity. **(A)** Mean layer-specific frequency-power decomposition of the BOLD signal in 21 ROIs (both cerebral hemispheres), averaged across 13 subjects. **(B)** Brain surface maps showing the average slope of the fitting line that best approaches the depth-varying ALFF per ROI. **(C)** ALFF slope for each cortical network. **(D)** The figure displays the regions that were included in the functional analysis, organized by network, over the surface of the brain (see **Table 1** for a detailed description of the ROIs included). **(E)** Brain surface maps showing the average ReHo slope of the fitting line that best approaches the depth-varying ReHo per ROI. **(F)** ReHo slope for each cortical network. **(G)** Representative ReHo map projected from six cortical surfaces to volume space. Results based on equi-distant layer sampling (for a comparison of ALFF and ReHo results in the whole-brain, gyri, and sulci when using an equi-distance or an equi-volume sampling model, see **Figures S7** and **S8**).

The assessment of layer-specific regional homogeneity (ReHo) demonstrated a more varied distribution (e.g., positive or negative slopes across ROIs), with a mean negative slope when considering the whole cerebrum (**Fig. S8**). **Fig. 2G** shows the ReHo map (mapped from surface to volume space) for a representative subject. The assessment of ReHo through gyri or sulci was conditioned by the model used to sample the layers along the cortical depth. An equi-distant layering resulted in curvature-specific differences, with a positive ReHo slope in most gyral locations (higher ReHo values in deep layers) and a steep negative slope in sulci (higher ReHo in superficial layers), both during rest and task (**Fig. S8A-B**). It is worth considering that, although large veins are more likely to traverse the outer portion of the brain (i.e. gyral locations), the presence of small venules over the sulcal surfaces could contaminate a larger number of nearby brain vertices, yielding higher homogeneity measures among superficial sulcal vertices. However, the differences in the ReHo slope between gyral and sulcal locations were not reproduced using an equi-volume sampling (**Fig. S8D-E**). The equi-volume sampling takes into account that the actual cortical folding leads to thicker superficial and thinner deeper laminae in sulci (and vice-versa in gyri). The heterogeneity of gyri/sulci ReHo results indicates that curvature-relevant assessments should be interpreted with caution and that the employed layer sampling model should be considered in this regard.

In addition to using ALFF and ReHo to assess brain activity, we investigated the functional connections between different brain regions at different depths of the cerebral cortex. To assess the reliability of our methods, we first conducted a layer-to-layer analysis within the visual system, following Polimeni et al (Polimeni, 2010). Briefly, we computed the temporal correlation, i.e. a proxy of functional connectivity, between the mean time courses of six different depths in the primary visual cortex (V1) of the right and left hemispheres, as well as between these and the layers in the secondary visual cortex (V2). In agreement with the literature (Polimeni, 2010), we observed that the superficial layers of V1 are strongly connected to other superficial layers of the contralateral V1, and that the superficial-intermediate layers of V1 connect mostly to intermediate-deep layers of the ipsilateral V2, probably reflecting a feedforward projection (**Fig. S9**). Next, we conducted a whole-brain functional connectivity analysis. This was done by generating a functional connectivity matrix from each fMRI scan, which included the temporal correlation between all pairs of time courses, i.e. between the average time course of each region at each layer and that of every other region and layer (i.e. 252^2^ cells). The correlation values of the six layers in each ROI and the six layers of every other ROI were computed. For group analysis, the connectivity matrices of 13 subjects were averaged and subjected to a t-test for statistical thresholding. The resulting adjacency matrix was then used to generate a layer-specific, whole-brain connectivity graph. A whole-brain connectivity map was computed for each of the three scans acquired per subject, and the results are shown in **Fig. S10I-L**. To facilitate the visual inspection of the depth-dependent connectivity graphs, the six assessed layers were grouped into “deep,” “intermediate,” and “superficial” layers, and connections were color-coded based on the pair of cortical depths involved, e.g., deep–deep connections were represented by yellow lines, intermediate–intermediate connections by red lines, deep-intermediate connections by orange lines, etc. In order to simplify the connectivity results, inter-regional connections were grouped per network. **Fig. 3A-D** shows four versions of four network connectivity graphs, with variable layer visibility, demonstrating the depth-specific significant connections within the DMN, the executive control network, the sensory-motor network, and the visual network, obtained from a resting-state scan acquired with protocol-1 (N=13). The same graphs were also computed from the task-based fMRI obtained with protocol-1 and the rs-fMRI obtained with protocol-2, which can be found in **Fig. S11**. A connectivity analysis based on coherence instead of temporal correlation rendered similar results (**Fig. S12**). The assessment of the cortical depth-dependent connectivity for each functional network, i.e. the average temporal correlation between ROIs of the network at each cortical depth, indicates that the superficial and upper-intermediate layers govern most resting-state correlations within the DMN and the visual network (bell shapes with a peak on the left in **Fig. 3F** -mean depth-dependent network connectivity across subjects (after transforming subject-specific measures to z-scores)-). The intermediate layers are seen to be strongly connected within the sensory-motor network (symmetric profile with a peak in the center in **Fig. 3F**), and it is the deep-intermediate portion of the cortex that exhibits the strongest connectivity within the executive control network (peak towards the right in **Fig. 3F**). The shift in laminar connectivity preference towards the infragranular layers in the executive control network was evidenced, for instance, as stronger interhemispheric connections between intermediate layers of the middle frontal gyrus compared to its superficial layers (significant connections in yellow and orange but not blue, between some ROIs in **Fig. 3**: “Executive control”). The laminar connectivity profile of each network computed for all subjects is presented in **Fig. 3E** (before z-score normalization). From a global-ROI perspective, i.e. independent of cortical depth, our data presented multiple significant connections that were in agreement with the volume-based resting-state network results derived from an ICA analysis; volume-based examples and the corresponding depth-dependent profiles per ROI are shown in **Fig. S13**.

**Fig. 3.**
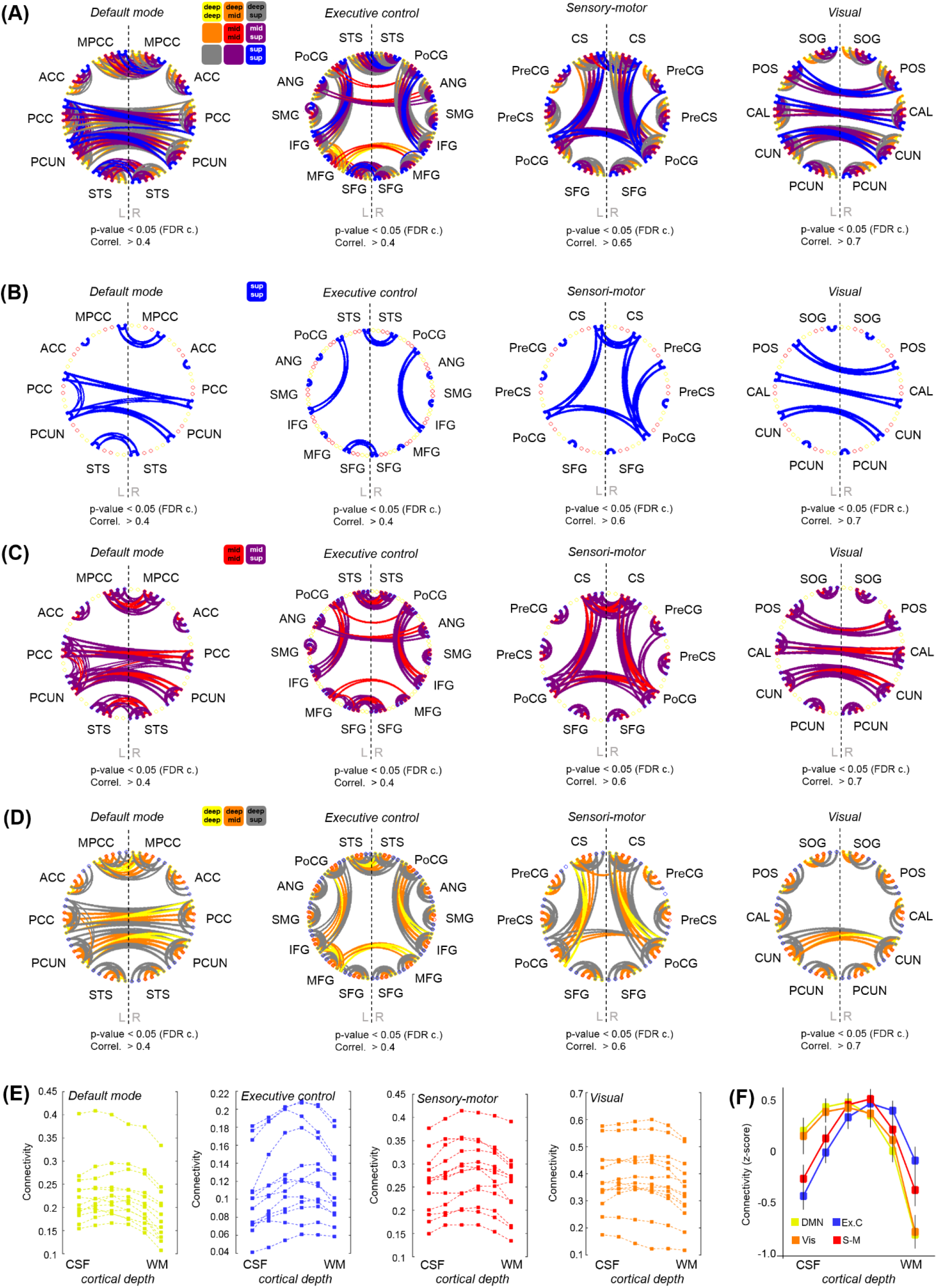
Cortical depth-dependent functional connectivity. **(A)** Four graphs showing the layer-specific functional connections between ROIs engaged in four different brain networks. A legend is provided above the first graph. A version of these graphs hiding the different layers is provided in **(B)**, **(C)**, and **(D)**. **(E)** Each of the four graphs shows the averaged connectivity (temporal correlation) within each functional network at each cortical depth for all subjects included in the analysis. **(F)** Cortical depth-specific global connectivity averaged in four resting-state networks (after transforming subject connectivity values to z-score). Error bars indicate ± standard error of the mean, N=13.

The identification of stronger connections involving the supragranular layers of the cortex in areas of the DMN is in agreement with previous reports demonstrating a higher level of activity and communication near the surface of the cortex in humans and non-human primates during rest (Guidi et al., 2020, Mishra et al., 2019).

### 3.5. State-dependent cortical-depth specificity of whole-brain functional measures

Having acquired whole-brain scans with high spatial specificity during rest and task performance, we aimed to assess the potential differences between both behavioral states at a laminar level. The slope or gradient of ALFF and ReHo measures along the cortical thickness, i.e. the difference between superficial and deep layers in terms of their signal amplitude and regional homogeneity, was significantly reduced when subjects were engaged in a motor task (**Fig. 4A** and **Fig. 4B**), suggesting that the superficial layers of the cortex are a major factor in the maintenance of resting-state activity that is significantly dampened during task performance. **Fig. S7C,F** and **Fig. S8C,F** show the differences in ALFF and ReHo laminar slopes, respectively, between resting-state and task using either an equi-distance or an equi-volume layering approach. In most cases, irespective of the sampling model, the laminar slopes were significantly higher during rest compared to task (i.e. higher ALFF and ReHo in the superficial layers).

**Fig. 4.**
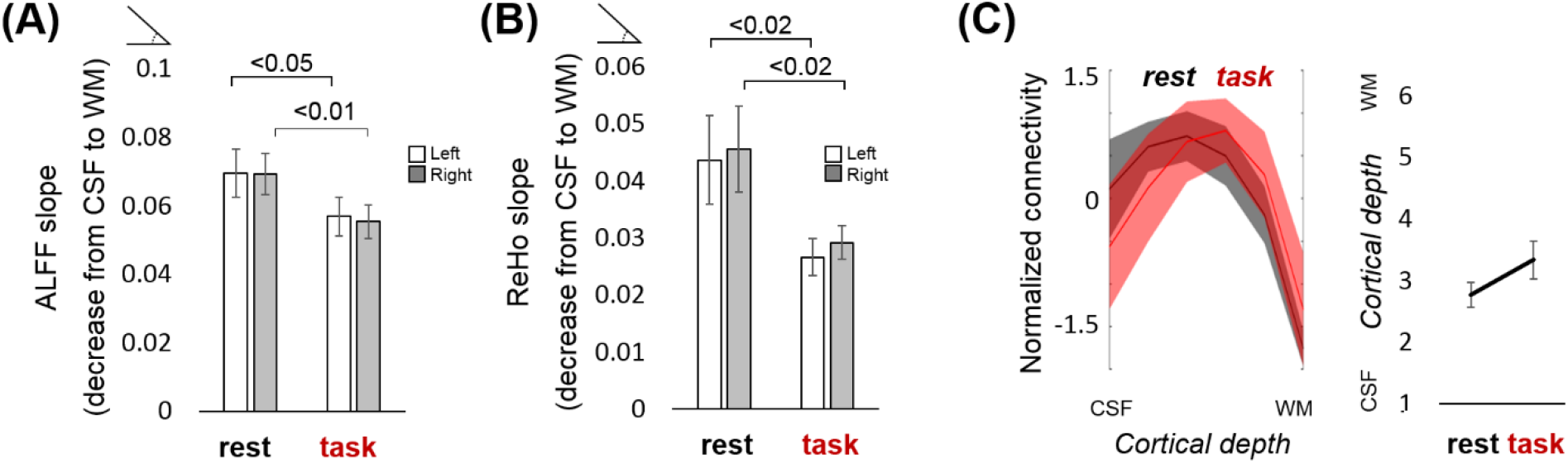
Functional laminar differences between resting state and task. **(A)** The bar plot represents the average ALFF slope computed for the resting-state and the task scan (after regressing out the estimated task response). **(B)** Average ReHo slope computed for the resting-state and the task scan (after regressing out the estimated task response). Values above the horizontal lines refer to the p-value of a t-test between conditions. **(C)** (left:) Normalized mean ± standard error of the laminar connectivity profile in the DMN during rest (black trace) and task (red trace), overlaid for comparison, and (right:) mean cortical depth at which the connectivity is maximum for each condition (not significantly different; the p-value of a t-test between conditions was 0.16). Error bars represent the standard error of the mean. N=13 for rs-fMRI-1 (“rest”); N=12 for task-fMRI (“task”).

Compared to task fMRI, resting-state scans generally exhibited higher connectivity between remote regions and their layers, as observed in their corresponding whole-brain connectivity graphs in **Fig. S10I-L** and in the network-specific graphs of **Fig. S11**. During the motor task, the connectivity within the executive control network seemed to involve more superficial layers of the cortex compared to rest (note the peak shift towards the left in the blue trace of **Fig. S10G or Fig. S10H** -with or without regression of the task paradigm-compared to **S10E**)). In contrast, more intermediate layers were involved in DMN connectivity during task performance (yellow trace in **Fig. S10G or Fig. S10H** compared to **Fig. S10E,** and **Fig. 4C**). This was observed, for instance, as a stronger correlation between the anterior cingulate cortices of both cerebral hemispheres and a reduced interhemispheric connection of superior temporal and posterior mid-cingulate areas (**Fig. S11A**, middle graph, compared to the left graph).

## 4. DISCUSSION

The layer-specific communication patterns across distant brain regions of the human brain have remained largely unexplored, mainly due to the limited brain coverage afforded by most high-resolution methods. In this report, we employed a recently developed fMRI sequence (TR-external EPIK) to address the question of how the different depths of the cerebral cortex participate in the maintenance of the resting state in humans. We first obtained motor-evoked responses that reproduced the laminar results seen in the existing literature and then provided a cortical-depth specific characterization of resting-state activity spanning multiple brain regions. Our ALFF showed a near-linear trend peaking at the surface of the cortex, likely reflecting vein-related bias, the slope of which was significantly reduced when subjects were engaged in a motor task. The slope of the mean ReHo across layers was also reduced during task performance, and network analysis showed certain differences both between networks and between conditions in terms of the cortical depth at which the connectivity was strongest, e.g., the DMN connectivity peaked at deeper layers in scans acquired during task performance. We interpreted the differences between resting-state data and task data as an indication that functional laminar dynamics can change with the brain state. Although our data may be partially contaminated by the influence of large veins (typical of GE sequences), certain metrics, especially those based on connectivity measures, demonstrated the sensitivity of the sequence to laminar events. Our results suggest that functional connections relevant to particular systems are cortical-depth specific, e.g., we found that intermediate and superficial cortical depths maintain key connectivity patterns in humans (e.g., DMN), in agreement with animal studies (Weiler et al., 2008, Whitesell et al., 2021), but we also identified a laminar connectivity shift towards deeper cortical layers in particular networks (e.g., executive control network).

In this study, we acquired two sets of rs-fMRI data for each subject within the same imaging session: one using isotropic voxels and the other using higher in-plane spatial resolution but with thicker slices. Both sets of data had voxels of similar volume (~ 0.25 mm^3^). The resemblance between all functional measures assessed in both sets of the rs-fMRI data (**Fig. S10** and **Fig. S11**) suggests that both protocols were able to detect brain connectivity with layer-specificity to a similar level of performance. Given the agreement between both protocols, and since protocol-2 confers a more complete brain coverage, despite the non-isotropic voxel dimensions in protocol-2 (0.51 × 0.51 × 1.00 mm), it may be a better candidate than protocol-1 for whole-brain laminar fMRI, especially in applications where structures that are located at the base of the brain, e.g., hippocampus, brainstem, etc., are significant for the integrity of the study, as long as the high-resolution plane matches the right dimension in the target area. A more detailed technical description of both of the TR-external EPIK protocols can be found in reference (Yun et al., 2022, Yun, 2022).

In previous literature, higher activation of superficial, i.e. supragranular, cortical layers has been observed in laminar rs-fMRI studies that have focused on a restricted FOV in the human brain (Guidi et al., 2020), in non-human primates (Mishra et al., 2019), and in rodents (Weiler et al., 2008). In contrast to the granular layer, which receives thalamo-cortical afferents, and the deeper, i.e. infragranular, layers, which mainly initiate efferent responses through subcortical and spinal projections, the superficial layers are the principal source and target of cortico-cortical regulatory communication (Moerel et al., 2019, D’Souza and Burkhalter, 2017, Rubio-Garrido et al., 2009, Larkum, 2013, Sempere-Ferrandez et al., 2019, Rolls and Mills, 2017). Hereby, the deeper portion of layer III receives long-distance feedforward projections, whereas its superficial portion, together with layer II, is targeted by long-distance feedback projections (Zilles and Catani, 2020). The present results support the supposition that superficial layers play a role in resting-state maintenance, as resting-state networks mostly involve cortical areas (Raichle et al., 2001, Heine et al., 2012); however, our results also demonstrate a network-specific laminar connectivity profile, i.e. not all networks’ connectivity peaked in intermediate/superficial territories. For instance, the connectivity within the sensory-motor network was stronger in the intermediate layers, which is consistent with feedforward connections from the thalamus to the sensory areas (e.g., postcentral gyrus), while higher-order networks, such as the DMN or the executive control network, mostly involved superficial or deep connections, suggesting feedback processing. However, it is worth noting that although directionality has been well studied within some sensory systems, e.g., in the visual pathway, the laminar preference of feedforward and feedback projections in other brain areas, and especially at the network level, is less well understood (for a review, see reference (Rockland, 2019)); hence, interpretations in this regard should be made with caution.

Although the literature upholds the results obtained from our resting-state analysis, i.e. strong participation of the superficial layers during rest (here observed with ALFF, ReHo, and connectivity measured within the DMN) or particular connectivity patterns in the visual system (e.g., feedforward connections between V1 and V2), the GE nature of the sequence employed poses the question as to whether the recorded signal could be merely a reflection of a vascular-related superficial bias, potentially further enhanced by a higher SNR near the surface of the brain (i.e. stronger T2 signal near the CSF). While the amplitude of the fMRI signal was generally higher in the superficial layers (measured as ALFF during rest and also indicated by the larger evoked responses on the superficial layers during task performance), likely partially due to the higher signal produced in superficial vascular territories, the varied connectivity results shown here demonstrate that the ability of the fMRI sequence to map depth-specific connections was not compromised to the extent of masking interactions occurring below the surface of the cortex. In fact, the connectivity measures rarely peaked on the most external layer of the cortex, and in some cases, the highest connectivity values were found in deeper territories; our results showed a network-dependent behavior of laminar connectivity. Besides, the fact that the layer-dependent signal amplitude, regional homogeneity, and whole-brain connectivity were significantly altered upon task performance using exactly the same imaging protocol (**Fig. 4**, **Fig. S10**, **Fig. S11**) suggests that a vein-related bias, if present, was not the only factor accounting for the observed responses. Notwithstanding this assertion, the potential implications of using a GE-based sequence will be discussed in a later paragraph.

The analysis of ALFF and ReHo in cortical regions of marked curvature (20% top and bottom curvature values, i.e. sulci or gyri) was conducted on six cortical surfaces extracted through equi-distant sampling and on six cortical territories segmented based on an equi-volume model. While both approaches provided similar results at a whole-brain level, gyri and sulci were characterized by different ReHo laminar profiles depending on the employed sampling method; in particular, the differences encountered when using the equi-distant sampling practically vanished with the equi-volume approach. This is probably due to the fact that the actual folding of cortical laminae leads to thicker superficial laminae in sulci vs. gyri (Bok, 1929), which is reproduced with an equi-volume but not with an equi-distant sampling (Waehnert et al., 2014). However, it is worth adding that our analysis based on equi-distance sampling was performed on surface-space, i.e. ReHo was conducted within each surface, while the ReHo results from equi-volume-based were obtained in voxel space first and later mapped to each territory in the segmented cortical ribbon. The latter approach is likely to be influenced by voxels from neighboring depths. In any case, our results suggest that, while depth-dependent levels of homogeneity evaluated in the gross cortex are similarly informed by both sampling approaches, and hence seem robust, finer differences pertaining to ReHo in sulci and gyri should be interpreted with caution.

In comparison to the resting-state paradigm, the assessment of layer-specific correlations during task performance revealed a decreased number of functional connections (**Fig. S10K** vs. **Fig. S10I**), which is in agreement with the literature (Jurkiewicz et al., 2018). The specificity of the responses evoked by the motor task was demonstrated with distinct line profiles in the precentral gyrus, in response to somatomotion, with and without somatosensation (**Fig. 1K**). Finger movement resulted in activation of the superficial and, to a lesser extent, deeper layers, the latter possibly indicating signaling from premotor and supplementary motor cortices, also coinciding with cortical motor efferents towards pontomedullary or thalamic nuclei (Weiler et al., 2008, Hooks et al., 2013). Sensory processing triggered by the touch between two fingers produced activation in intermediate layers, possibly reflecting the modulation of the motor cortex by sensory cortical areas and afferents from the sensory thalamus (Weiler et al., 2008, Hooks et al., 2013, Mao et al., 2011). The activation observed near the cortical surface may reflect, at least to a certain extent, a contribution from the venous network; however, our laminar profiles do not merely present a linear vein bias. Moreover, superficial neuronal responses can be expected during the performance of a motor task, as motor thalamic afferents modulate motor behavior in the superficial and intermediate layers of the cortex in a similar way to the sensory thalamic afferents upon the addition of touch (Hooks et al., 2013) - which has also been observed in laminar studies (Huber et al., 2017). The fact that evoked responses in the intermediate and deep layers were detected in a line-profile analysis but not in the analysis of a bigger cortical patch, also responding to the motor task but exhibiting an upper → deeper trend only, demonstrates the specificity of the laminar responses on particular cortical regions and the overall preponderance of BOLD responses in the superficial layers in less specific evaluations. Although some corrections could be applied to the evoked responses to compensate for the vasculature-related gradient, we deferred from using this procedure. This was done to avoid masking the contribution of non-neuronal sources that would be present in the resting state data, i.e. we opted for applying the same correction to task and resting-state data, as correction of the latter using existing models is currently not straightforward.

It could be argued that, aside from the potential venous influence, the higher SNR in the superficial layers (e.g., **Fig. 1E**, upper layers showing higher mean MR signal in the TR-external EPIK image) could facilitate the detection of functional responses and connections between superficial cortical depths. However, as superficial, intermediate, and deep connections pertaining to different network systems were detected differently during task performance and during rest (e.g., **Fig. 4C**, **Fig. S10**, **Fig. S11**), and the ALFF and ReHo laminar gradients were reduced during task performance (**Fig. 4A-B**), it is likely that functional information remains in the high-resolution data, despite potential bias in favor of the superficial layers; i.e. not all activations and connections were observed in the most external layer of the cortex or to the same extent, independent of the behavioral condition. The evaluation of typical resting-state measures in our task-fMRI scans after regression of model activation responses provided a reference framework of potential changes at the level of cortical laminae relevant to the brain state. Although most connections that are typically found during the resting state exist in task-related data, our results showed that the preferred layers to maintain such connections can change, e.g., the DMN during task appears less supported by superficial layers than during rest (**Fig. 4C**, **Fig. S10G vs. Fig. S10E** and **Fig. S11A**). It is well known that the connectivity of the DMN decreases during task performance (Raichle et al., 2001), which is consistent with the obtained results and may also explain the shift from superficial to intermediate layers (possibly reflecting reduced feedback processing during task performance); however, a decrease of activity in DMN areas is also expected to be bound to a reduced blood flow and, hence, to a potential improvement (reduction) of the superficial bias, which should be taken into consideration when interpreting the task-related whole-brain results.

The selected fMRI sequence integrates a number of acceleration techniques that enable a large matrix with fine resolution to be sampled without compromising the quality of the data (the sequence implementation has been previously demonstrated in comparison to EPI (Yun and Shah, 2017, Yun et al., 2013)). In TR-external EPIK, the high spatial resolution is partially possible due to the segmented sampling of the k-space periphery. While full k-space sampling is completed every 3 TRs, the combination of this with a fully sampled center k-space at each TR confers upon TR-external EPIK enough temporal resolution to sample brain hemodynamics with sub-millimeter resolution ((Yun et al., 2019, Yun and Shah, 2017, Yun et al., 2013, Yun and Shah, 2019, Yun, 2022)). The results from the group of subjects studied in this work demonstrate the feasibility of using TR-external EPIK for the detection of layer-specific functional signals. In our protocol, the sharing of missing k-space lines is limited to the adjacent two scans, the energy of which was determined to be smaller than 10% of the entire k-space energy. Although the peripheral k-space is updated slowly (i.e. 3 TRs = 10.5 s; ~0.1 Hz), it is sufficient to track the change of resting-state signals, the peaks of which are usually detected within a frequency of 0.03-0.04 Hz (Zuo et al., 2010, Xue et al., 2014, Yuen et al., 2019, Bajaj et al., 2014) in the gray matter. Furthermore, it has been reported that conventional resting-state analyses are not significantly altered by TR (Huotari et al., 2019), and methods employing relatively slow sampling have succeeded in tracking laminar dynamics (e.g., 3960ms (Sharoh et al., 2019), 5000ms (Barry et al., 2021), 8300ms (Huber et al., 2020)). Despite the advantages of TR-external EPIK over conventional EPI, the GE nature of the technique makes it susceptible to vein-derived contamination, just as any other GE-EPI sequence. Assuming a fair neuro-vascular coupling, the fMRI signal extracted from a cortical capillary network is a reliable indicator of neuronal activity. However, capillaries supplying blood to neurons flow in bigger ascending venules that run perpendicular to the cortex and disgorge into pial veins tangential to the surface of the brain, and these veins influence the fMRI signal and can result in a bias towards the superficial layers of the cortex. This bias is reduced in other acquisition schemes, such as SE-BOLD fMRI, sensitive to oxygenation changes in the microvasculature only, vascular space occupancy fMRI (VASO), sensitive to vessel dilation, or cerebral blood flow-based fMRI (CBF), sensitive to flow changes. Due to the short T2 constant of blood at ultra-high field, BOLD-fMRI at 7T is only sensitive to the extra-vascular effect of the oxy/deoxy-hemoglobin ratio, irrespective of vessels size. Although the parenchymal vessels, i.e. venules of diameter < 25 μm, are detected by both SE- and GE-BOLD, these are often overshadowed in GE sequences by the influence of bigger veins running along the surface of the brain, which cause static perturbations in the magnetic field affecting the neighboring parenchymal tissue. Therefore, while SE-BOLD offers a fine localization of the activated parenchyma, GE-BOLD provides a mixed signal that is relevant to both small parenchymal vessels and macroscopic veins. To correct for this bias, we subtracted the time-varying phase component of the MR signal from each voxel’s magnitude time course. The time-varying phase component is proportional to the frequency offset exhibited by the spins and hence, sensitive to the venous vasculature (Menon, 2002, Curtis et al., 2014). As large vessels produce field inhomogeneities, spins processing in the neighboring parenchyma would produce changes in the phase component of the signal; in contrast, small randomly oriented vessels, e.g., in the capillary network within the gray matter, do not produce a net phase and hence, their derived signal remains intact. Although this method helps to improve the local specificity of BOLD, several biases could remain in the corrected data, such as those derived from motion correction or geometric distortions (Bause et al., 2019), which reduce the ultimate spatial resolution of GE sequences. However, it is worth noting that geometric distortions are less pronounced in EPIK compared to conventional EPI, and in fact, the PSF of GE-EPIK is narrower than in conventional GE-EPI (Yun et al., 2013). A few groups have demonstrated the dependence of laminar fMRI on the orientation of the cortex with respect to B0 (Viessmann et al., 2019, Baez-Yanez et al., 2017). Although orientation bias can be critical for the dissection of the fMRI signal, this effect has been correlated with low-frequency drift, motion, and respiratory cycles, which were accounted for in our study, therefore diminishing the discussed effect. Moreover, most of our results were obtained from relatively big patches of the cortex and through multiple ROIs, averaging vertices with different orientations within a cortical surface. Hence, although non-negligible, an orientation bias is not expected to have conditioned our results. However, a future model accounting for such an effect could determine the extent to which the curvature affects the laminar profiles at a whole-brain level.

Although the 2D acquisition scheme employed in our TR-external EPIK sequence could raise concerns relating to the increased susceptibility to inflow effects compared to 3D-EPI, this increased susceptibility is only significant when a relatively short TR (i.e. tens of a millisecond) is used (Frahm et al., 1994). Hence, signal changes induced by the inflow effect are negligible in most 2D multi-slice fMRI protocols configured with a TR larger than 1.5 s (Speck and Hennig, 1998, Howseman et al., 1999, Wiggins, 2000, Gao and Liu, 2012). Since our protocol also uses a relatively long TR (3.5 s), the inflow effect on the detection of the fMRI signal can be considered insignificant. In EPIK, the multi-shot scheme for the peripheral k-space can lead to an increased sensitivity to the subject motion when compared to EPI. However, the sampling of the central k-space region at every temporal scan in EPIK can effectively ensure comparable temporal stability to the EPI case, enabling a robust acquisition of fMRI time-series data (Yun and Shah, 2017, Yun et al., 2013). As 3D EPI often requires additional correction methods to account for intra-scan motion artifacts, 2D EPIK can be more straightforwardly deployed in an fMRI study.

In contrast to retrospective model-based approaches that reduce the superficial BOLD bias by applying temporal decomposition (Kay et al., 2020), linear detrending (Fracasso et al., 2018), normalization (Kashyap et al., 2018a), or deconvolution (Heinzle et al., 2016, Markuerkiaga et al., 2016) to evoked fMRI data, the phase-based correction method employed here does not involve a priori assumptions about the behavior of the vascular dynamics across the cortical thickness and does not rely on the use of an evoked paradigm, offering a bias-free alternative to correct resting-state GE-fMRI data. Based on our results, this phase-based method (Menon, 2002) may constitute a rather conservative approach, i.e. the activity of the superficial layers was found to predominate almost ubiquitously (higher ALFF in upper layers of the cortex). However, connectivity analysis did identify laminar interactions in a cortical profile different from the linear gradient that would be expected in purely macro-vascular measures, i.e. with a maximum on the surface, and the evoked responses were composed of two peaks, the latter located in deeper layers of the cortex, indicating a decent laminar sensitivity. Future studies employing GE TR-external EPIK and SE TR-external EPIK in a combined acquisition will investigate the possibility of complementing the phase-correction method with additional strategies to exploit the high SNR of GE schemes with robust laminar specificity.

The vascular dependence of the fMRI signal remains one of the biggest challenges in laminar fMRI (Uludag and Blinder, 2018). Signals obtained with BOLD contrast are not only conditioned by a physically-constrained laminar point-spread function, i.e. the fact that ascending venules spread the deep activation signals towards superficial layers (Havlicek and Uludag, 2020, Markuerkiaga et al., 2016, Marquardt et al., 2018), but also by the varying physiological mechanisms underlying the BOLD response (CBF, CMRO2, CBV), which can modulate this point-spread function across cortical layers (Havlicek and Uludag, 2020). Despite the complexity of the model, laminar fMRI studies have even employed strategies that artificially sample the cortical depth with a number of points higher than the number of voxels across the cortical ribbon. This is appropriate under the assumption that the purpose of the study is to track the trend of activation across the cortical depth and not to resolve the actual neuronal signal at any particular layer, which is indeed not possible, even with sufficient spatial resolution, given the facts discussed above. In this study, the number of depths used was equal to the number of histologically-defined layers in the neocortex; however, the layers in our model have been arbitrarily chosen to be equally spaced along the cortical thickness and have no direct relationship with the cortical cytoarchitecture. Hence, our conclusions should be interpreted in terms of cortical depth only. Although some reports have chosen a smaller number of layers for their laminar assessment, e.g., three (Sharoh et al., 2019), which simplifies the analysis and confers layers with a greater independence, others have shown that a greater number of depths, e.g., twenty (Huber et al., 2017), enables clearer detection of cortical responses, despite the fact that this strategy involves a greater dependence on neighbor signals. In addition, the latter scheme also poses an important challenge in whole-brain studies due to the high computational demands of the analysis procedure (see discussion below).

Irrespective of the number of layers, ambitious whole-brain laminar fMRI studies should ensure perfect co-registration between the anatomical reference scan and the functional data. Previous reports have shown the possibility of integrating T1 contrast within a high-resolution fMRI sequence (e.g., (Huber et al., 2017), acquired during VASO), which allows a 1:1 relationship between the anatomical and functional scans, mitigating any co-registration concern. This option was not yet available in the current study. Nevertheless, in this work, we achieved acceptable co-registration in the majority of the brain by using manual rigid transformations (**Fig. S14**), which proved to be more reliable than the transformations rendered by non-linear automatic registration tools. In addition, the improved robustness against geometric distortions in TR-external EPIK (Yun et al., 2013) is expected to yield fewer co-registration errors, compared to the common EPI, between the functional scans and the anatomical scans (e.g., MR2RAGE). However, particular areas in different subjects remained poorly co-registered and needed to be masked out to ensure accurate laminar analysis. In the future, TR-external EPIK will be modified to provide a T1 contrast, which will avoid data loss, reduce the pre-processing time, and, more importantly, will alleviate the potential concerns relating to co-registration.

In this work, we employed TR-external EPIK to map the majority of the brain with a voxel resolution of 0.63 × 0.63 × 0.63 mm^3^ or 0.51 × 0.51 × 1.00 mm^3^. In addition to the challenge in terms of sequence design, the resulting matrix size imaged by TR-external EPIK (336 × 336 × 123 or 408 × 408 × 108) presents important computational limitations in post-processing, such as the relatively large memory required (minimum storage space: ~ 6 Gb per 4D-volume; RAM: ≥ 32Gb for many pre-processing steps) and long processing times. After pre-processing, analysis of the cortical surfaces with numerical computing environments such as Matlab is, again, conditioned by the number of vertices to be processed. Therefore, the greater the number of layers, the higher the computational demands. For instance, one cortical surface covered by protocol-1 contained ~ 290,000 vertices, which means that the six surfaces obtained from both hemispheres involved the analysis of ~ 3,480,000 vertices, each of them consisting of a 168-point time course. In this work, in order to adjust the computational demands to those of a standard scientific workstation, the first step in most of the layer-specific functional analyses was to average all vertices specific to one layer within one ROI. Given that the laminar performance of the cortex can be highly localized, some events might be overlooked when using an ROI-based analysis to identify particular behaviors; hence, the use of supercomputing resources to assess vertex-specific measures becomes highly desirable to exploit the information contained in the acquired data.

This study adopted a sequence offering high spatial resolution (~ 0.25 mm^3^ voxels) to map laminar functional dynamics in a 7T scanner with near whole-brain mapping in healthy subjects. We investigated the dependence of common rs-fMRI measures such as ALFF, ReHo, and functional connectivity on cortical depth, which has emerged as a new dimension to be explored in the human brain. Our results indicate that the cortical depth is differentially (functionally) connected depending on the resting-state network, with intermediate and superficial layers playing a critical role in the maintenance of the DMN, and further show that the laminar involvement can change during a motor task. This work demonstrates the potential of laminar analysis of connectivity in the study of broad cortical networks, which may have direct implications in the research of psychiatric disorders that are commonly associated with network disturbances.

## Acknowledgments

*We thank Dr. Michael Schwerter for guidance in the phase-signal processing; Ms. Elke Bechholz and Ms. Anita Köth for technical support; Ms. Claire Rick for manuscript corrections; Prof. Zuo for guidance on the usage of CCS, and the fMRI volunteers for their excellent cooperation.*

## Notes

**Conflict of Interest** *The authors declare that the research was conducted in the absence of any commercial or financial relationships that could be construed as a potential conflict of interest*.

### Competing Interest Statement

The authors have declared no competing interest.

### Summary of Updates

Based on the feedback obtained in recent revisions, we have updated some of the figures and added some explanations.

