## Supplementary Material for "Cortical depth-dependent human fMRI of resting-state networks using EPIK"

#### 1 Supplementary Data

**SUPPLEMENTARY METHODS:**

**Anatomical pre-processing**

The MP2RAGE volume was first co-registered to each functional image for each subject and processed with an automatic pipeline that extracts the inner and outer surfaces of the cortex (Freesurfer: *recon-all* + -hires) (**Fig. S1a**). The boundary between the cortical gray matter (GM) and the outer cerebrospinal fluid (CSF) is referred to as "pial surface". This is due to the proximity of this cortical surface to the pial matter, the innermost meningeal membrane surrounding the brain. The boundary between the GM and the white matter (WM) is referred to as "white surface", due to its proximity to the white matter. During the tessellation process (transformation of volume space to surface space), each voxel edge was converted into three vertices. The resulting pial and white surfaces were then smoothed by a factor of two (FreeSurfer: *mris_smooth*). To sample the cortical ribbon through its depth, four intermediate surfaces were generated by expanding the inner ("white") surface outwards by 20% of the cortical thickness (Freesurfer: *mris_expand –following an equi-distance model*). The four layers defined between the white surface and the pial surface are referred to as +20%, +40%, +60%, and +80%, respectively (+20% is the 2$^{nd}$ deepest layer, and 80% is the 2$^{nd}$ most superficial layer). Note that the word "layer" here does not refer to the cytoarchitectonically-defined cortical layers but instead to each of the cortical depths that are analyzed in our study. Given that the cortical thickness ranges from ~ 2 to ~ 3 mm, the average distance between consecutive layers varies between 0.4 and 0.6 mm, which is similar to the resolution of the employed imaging protocol. A discussion about the number of layers and fMRI resolution is provided in the *Discussion* in the main manuscript.

**Analysis of the anatomical data**

The cortical thickness was automatically calculated by the function *recon-all* at each vertex. The average thickness per ROI was calculated as the average thickness across all the vertices in an ROI. The surface curvature was used to distinguish between vertices forming a gyrus or a sulcus (1/radius, calculated by recon-all -see example in **Fig. S2a**-). A vertex was said to constitute a gyrus if its curvature value lay within the bottom 20% (negative curvature) of all the curvature values in the brain and was determined to be a sulcus if its curvature was within the top 20% (positive curvature). The average thickness was then calculated for all gyral and sulcal vertices independently. To characterize the surface model for sulcal and gyral vertices, a measure of the distance between adjacent vertices was calculated per layer (**Fig. S2b**), which can be interpreted as the in-surface resolution. Similarly, the inter-layer distance was calculated in gyral and sulcal locations (**Fig. S2d**) to provide an estimate of the signal dependence to be expected between adjacent layers on gyri or sulci. Most of the anatomical analysis was performed following Kay et al. [1]. In order to reduce the computational demands of the whole-cortex functional analysis, surface decimation, e.g., an extension of the number of vertices in each cortical surface, was not performed.

**Functional pre-processing**

Magnitude and phase images were reconstructed from each functional scan. The phase images acquired from our scans did not show any noticeable temporal or spatial wrapping (they were comparable to the result of applying a 4-D unwrapping algorithm) and were hence used in their raw format. After removing the first four volumes to ensure signal stabilization, the following steps were applied to both the magnitude and phase images: slice-timing correction (Matlab: SPM), realignment (performed first on the magnitude image and applying the resulting matrix to the phase image, AFNI: *3dVolreg/3dAllineate*), temporal filtering (0.005 - 0.12 Hz), regression of the physiological parameters (AFNI: *3dDeconvolve* -regressor estimation explained below-), motion regression (AFNI: *3dDeconvolve*) and regression of the mean white matter and CSF signal. The mean white matter and CSF time course were generated by averaging the time course of voxels within white-matter and CSF masks created from the mean fMRI volume. All the regressions and temporal filtering were applied in a single step using AFNI. The physiological regressors were generated by estimating up to the 5$^{th}$ order Fourier sine and cosine series fit of the cardiac and respiratory signals, plus multiplicative terms (Matlab: https://github.com/tesswallace/retroicor /blob/master/mod_retroicor.m). Subjects were screened and excluded from the analysis if mean head displacements exceeded 0.5 mm (this was true for one of the 18 subjects that were initially measured). The mean head displacement was calculated as:

$$mean(\sqrt{(diff(M_{r1}))^2 + (diff(M_{r2}))^2 + (diff(M_{r3}))^2 + (diff(M_{t1}))^2 + (diff(M_{t2}))^2 + (diff(M_{t3}))^2})$$

with $M_{r1..3}$= 3 rotational, and $M_{t1..3}$= 3 translational vectors of length= scan length (result from the realignment step).

Following Menon et al. [2, 3], a phase-based correction method to reduce the effect of veins on the GE-BOLD magnitude signal was implemented as follows. First, the coefficient "Fit" in a least-squares fitting of the pre-processed phase time course ($\phi$) to the pre-processed magnitude time course (M) was calculated in a voxel-wise manner, such that M = Fit$^{\phi}$ + residuals, where the sum of squared residuals is minimized (AFNI: *3dTfitter*). Fit$^{\phi}$ represents the signal in M that can be explained by $\phi$ and is attributed to the effect of field inhomogeneities, e.g., contributions of large veins. Fit$^{\phi}$ was then subtracted from M (AFNI: *3dcalc*). The procedure achieves signal correction in venous voxels while leaving unaltered the signal of voxels in the gray matter (**Fig. S3**).

Due to the need for perfect co-registration between the anatomical and the functional datasets, the mean magnitude image of each subject was manually aligned (with linear transformations and cubic interpolation) to its corresponding MP2RAGE using Freeview, and the inverse registration matrix of this operation was applied to the anatomical image before anatomical pre-processing to produce cortical surfaces in the functional image space directly. To avoid erroneous mapping of functional voxels to the MP2RAGE-based cortical surfaces, a mask was drawn for each scan and for each subject over the areas that remained poorly aligned after manual co-registration ("error-mask"), and voxels within the error-mask were removed, i.e., not projected to the cortical surfaces. This ensured near-perfect co-registration between MP2RAGE and TR-external EPIK on the portion of the cortical ribbon involved in the subsequent analysis (**Fig. S14**). The corrected functional image was then mapped onto the six cortical surfaces to generate functional layers (Freesurfer: *mri_vol2surf*). Smoothing up to 1mm was applied in surface space, i.e., within each cortical layer independently, using the flag “surf-fwhm 1” in *mri_vol2surf*.

**Task-related analysis**

Functional volumes were subjected to a General Linear Model (AFNI: *3dDeconvolve*), and the resulting beta maps were mapped to the cortical surfaces and smoothed within each layer. (Freesurfer: *mris_smooth*). After layer-specific smoothing, surface beta maps were projected back to the cortical ribbon in volume space (Freesurfer: mri_surf2vol).

*ROI-based results*

For ROI-based evaluation of the evoked responses on the primary motor cortex, a cortical patch of a region of the left pre-central gyrus was first extracted and flattened (Freesurfer: *mri_annotation2label, label2patch, mris_flatten*). Then, the six surface beta-maps for each subject were overlaid on the cortical patch, and the mean beta value was computed at each cortical depth using Matlab. For each subject, the beta profile was normalized to the maximum beta value along the cortical thickness. The mean ±SD among 13 subjects was calculated (**Fig. 1b**).

*Line-profiles*

The surface beta-maps projected back to the cortical ribbon were subjected to a line-profile analysis (underlay: cortical segmentation, overlay: beta-map). A quadrilateral shape was defined, covering a small portion of the lateral side of the hand knob (area M1-4a), extending from the CSF towards WM (**Fig. 1d, f or h**), and 20 lines were generated to sample the beta map along the cortical thickness, with each line sampling 100 points. The 100-point profiles of the 20 lines were averaged, and the mean ±SD was plotted (**Fig. 1i**). The cortical limits (CSF-GM and GM-WM) were identified by generating the line profile of the cortical segmentation image (**Fig. 1f-g**). A customized Matlab script was used to load the particular slices and generate the line-profiles. After normalization, which consisted of dividing the line profile of each subject by its maximum value, the line profiles of 13 subjects were averaged to show mean beta ±SD (**Fig. 1c**).

**Resting-state analysis**

*Volume-based functional analysis of resting-state networks*

The corrected functional volumes were subjected to an independent component analysis (FSL: *melodic*). Eighty components were extracted, and networks were identified by visual inspection of the probability maps thresholded at 0.5. In the current study, these maps were used to confirm the presence of resting-state networks in the data with a standard method in volume-space.

*Layer-specific functional analysis: ROI definition*

Twenty-one ROIs were selected from a surface atlas (Freesurfer lh/rh.aparc.a2009s.annot, based on the Destrieux atlas -see **Table 1**-). The functional surfaces were loaded in Matlab (Matlab: *MRIread*) and the functional time courses of vertices belonging to each ROI and cortical depth were averaged to reduce the data dimensionality. In total, 21 average time courses were extracted per layer in each cerebral hemisphere.

*Layer-specific frequency-power decomposition*

A frequency-power decomposition was obtained from each functional time course for frequencies in the range 0.01-0.1Hz in bins spanning 0.002 Hz (Matlab: *periodogram*). The plots in **Fig. 2a and Fig. S10a-d** are the result of averaging the power of the fMRI signal per frequency bin and per cortical layer in each of the 21 ROIs across 13 subjects.

*Layer-specific ALFF and ReHo*

To quantify the amount of activity in a particular ROI, the amplitude of the low-frequency fluctuations (ALFF) was calculated. This measure was computed voxel-wise in volume space (AFNI: *3dRSFC*, 0.01 – 0.1 Hz), converted to z-score (by subtracting from each voxel the brain mean and dividing by the standard deviation), and was then mapped to surface space (six ALFF surface maps per hemisphere). Values were averaged across vertices for each ROI, and the mean ±SD of 13 subjects was calculated. Regional homogeneity (ReHo) measures were calculated in two ways: directly in surface space (per-vertex calculation) considering two neighbors using a matlab function adapted from CCS (Connectome Computation System [4]), or in volume space (AFNI: *3dReHo,* selecting 19 neighbors), with the voxel values being mapped to the cortical surfaces later on. The surface analysis was used to generate the results shown in the main figures. These were derived from an equi-distant sampling and surface generation on Freesurfer. To evaluate the effect of the sampling method on ReHo and ALFF, in addition to the equi-distant sampling from Freesurfer, an equi-volume sampling was produced using LAYNII (https://github.com/layerfMRI/LAYNII, [5]), which yields cortical segmentation in voxel space. The ReHo values were later normalized to z-scores (by subtracting the mean of all vertices across all surfaces and dividing by the standard deviation) and averaged per ROI. In both cases, three surface ROI analyses were conducted in parallel: a) considering all the vertices of each ROI; b) considering only the vertices on the 20% top gyri (sorted by curvature); c) considering only the vertices on the 20% top sulci. Mean values were plotted per cortical depth for all three strategies. Data from the two resting-state scans and the task scan (full time courses after pre-processing) were analyzed for comparison.

The dominance of the superficial layers was further evaluated by assessing the rate at which any particular measure changed with respect to the cortical depth. Given the quasi-linear behavior of the laminar ALFF and ReHo profiles, we performed a linear fitting using Matlab (*polyfit*), and the slope (linear coefficient) of the fitting line was stored for each ROI and subject. The slopes of fitting lines corresponding to ROIs belonging to a functional network were averaged, and the mean slopes were averaged across subjects. A paired t-test was computed for each functional network between the measures obtained in sulci and those obtained in gyri to detect significantly different slopes in cortical areas of different curvature. In addition, a t-test considering whole-ROIs, sulci, and gyri was conducted between the slope values obtained during resting-state or during task performance. P-values below 0.05 were considered significant.

*Layer-specific functional connectivity*

Functional connectivity was calculated as the correlation coefficient (Matlab: *corr2*) between each pair of functional time courses (six functional time courses per ROI and hemisphere). A t-test was computed for all connectivity matrices in a group (N=13). Connectivity graphs were plotted as lines connecting layers in ROIs exhibiting a thresholded statistic and correlation value (specific values of thresholding shown in figures). All connectivity results related to the resting-state networks are based on temporal-correlation analysis unless reported otherwise (e.g., for a comparison, coherence analysis was performed between the ROIs of four functional networks, as shown in **Fig. S12**). Global connectivity measures were normalized within each subject and network by dividing the correlation value between each pair of layers by the mean correlation value in the layer connectivity matrix.

In addition to the cross-laminar/ROI correlation analysis, the representative time course of each independent component associated with a functional network in volume space was used to identify the depth-dependent correlates of each functional network. For each scan of each subject, a volume-based independent component analysis (ICA) was run using FSL (*melodic*), which resulted in eighty independent components, each of them characterized by a temporal time course. Four independent components (ICs) relevant to four functional networks (default mode, executive control, visual, and sensory-motor) were manually selected based on their corresponding spatial maps. The correlation coefficient between each IC and the time course of all vertices in each functional surface was calculated, and the correlation values of vertices within each ROI and each cortical depth were averaged. This resulted in a layer-ROI correlation matrix for each IC and each subject. Correlation matrices corresponding to each particular IC were averaged across subjects.

***Resting-state measures taken from the task-fMRI data***

To examine brain behavior with laminar-specificity during task performance, which could serve as a reference of a different brain state compared to the resting-state fMRI, the same analysis as in the resting-state data (ALFF, ReHo, connectivity) was applied to the task fMRI data after regressing out the task predictor time course (convolution of the stimulation paradigm with the HRF –as in AFNI 3dDeconvolve defaults-) from fMRI voxels.

***Statistical analysis***

To investigate the significance of the results, a t-test was performed in ALFF, ReHo, and connectivity analysis. Due to the large number of samples (6 layers per ROI, 42 ROIs), the threshold p-value for significance in the ROI-connectivity analysis was corrected using a false discovery rate control method, as implemented by the matlab function fdr_bh.

**Vascular occupancy analysis**

The T2/T2*-weighted mean surface, obtained as an average of the realigned volumes co-registered to MP2RAGE and mapped to each cortical surface, was used to calculate the number of venous vertices per cortical depth. Venous vertices were identified as those exhibiting an intensity value below one standard deviation in the range of image intensities in a given scan (the mean standard deviation of the six layers was used as the threshold). This signal threshold corresponded to an approximate value of 300 in the mean image obtained at 7T. A histogram of the average intensity values extracted from the whole brain across the six different layers is shown in **Fig. S5a**. An example of the venous surface maps for the pre-central gyrus in a representative subject is displayed in **Fig. S5b**.

### 2 Supplementary Figures

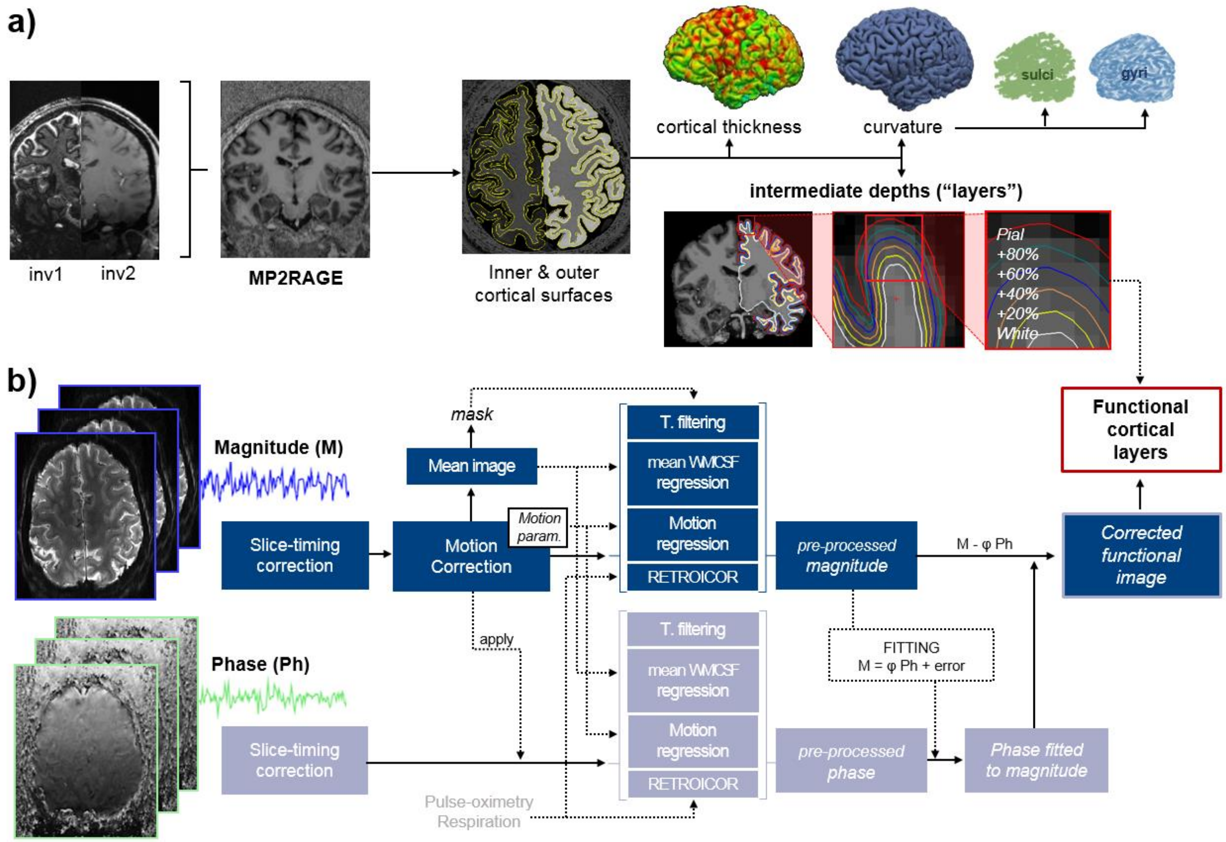


**Figure S1. Main steps for the pre-processing of the anatomical (a) and functional scans (b). a)** "inv1" and "inv2" indicate the first and second inversion contrast images of MP2RAGE. The T1-weighted MP2RAGE is used to segment the brain into gray matter, white matter, and cerebrospinal fluid (CSF). The cerebral cortex (outer brain gray matter) is further processed to extract its boundaries (outer or "pial" and inner or "white"). Several calculations can be performed based on these surfaces (e.g., thickness, curvature). Four additional surfaces are generated at equal spacing. **b)** Both the magnitude and phase images obtained with TR-external EPIK are subjected to a routine pre-processing pipeline, including slice timing correction, motion correction, regression of physiological signals, and temporal filtering. The pre-processed phase image is used to correct the magnitude image from vein-related bias. The corrected pre-processed magnitude functional image is then projected onto the six generated surfaces, in subject space, for laminar fMRI analysis.

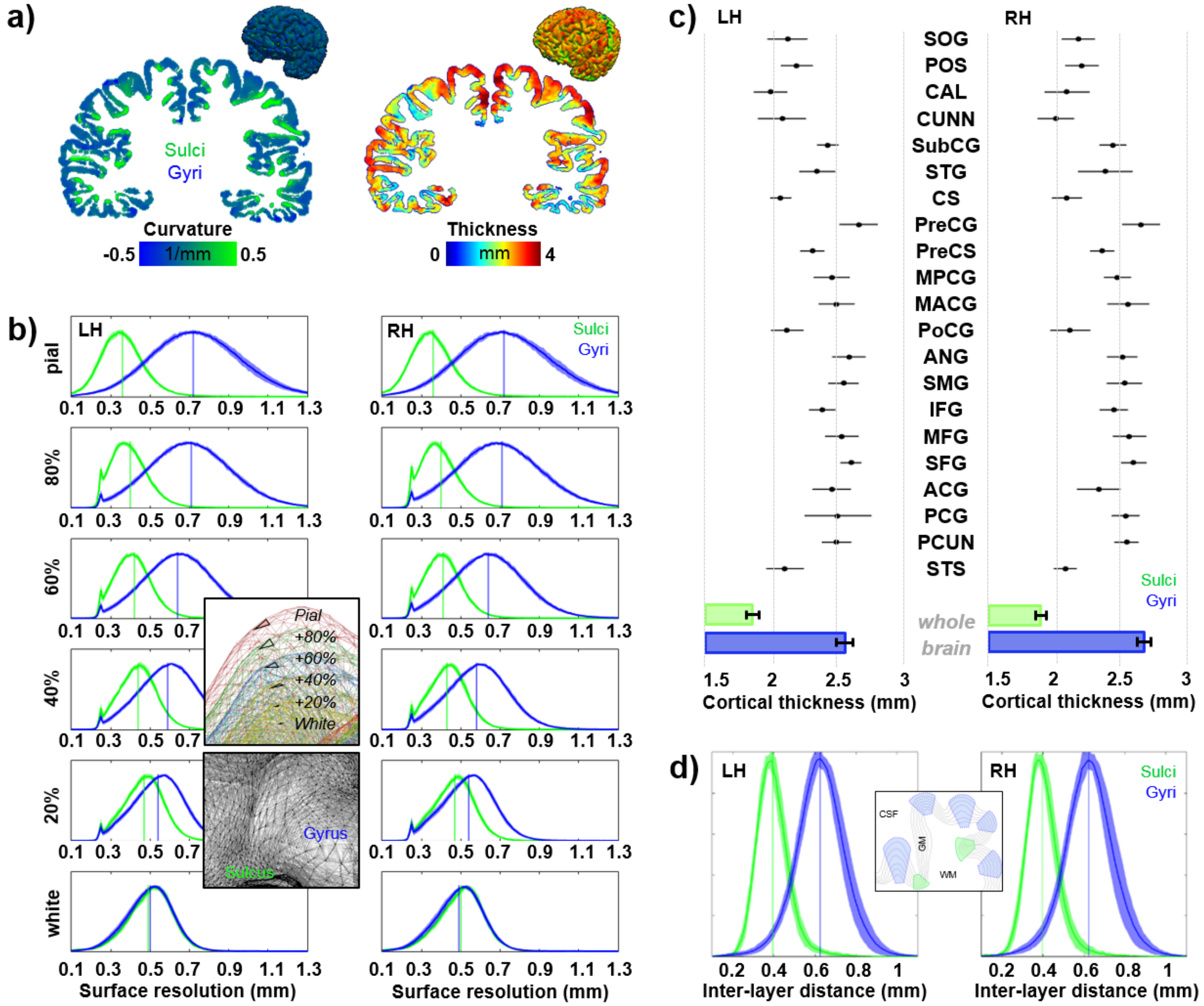


**Figure S2. Evaluation of the cortical model. a)** Coronal slices showing the curvature and thickness of the cerebral cortex in a representative subject. **b)** Histograms showing the distribution of distances between two consecutive vertices at six different cortical depths for locations corresponding to gyri (blue traces) or sulci (green traces) in both cerebral hemispheres. The insets show an example of the cortical surfaces in a gyrus (upper inset –an individual triangle is highlighted on each surface-) and of the pial surface over a sulcus and a gyrus (lower inset). **c)** Cortical thickness calculated in 21 ROIs and mean cortical thickness calculated from the whole brain in gyri and sulci for both cerebral hemispheres (mean ±SD, N=18). **d)** Histograms showing the distribution of the distance between vertices located in contiguous layers (e.g., white–layer02, layer02-layer04; etc.), for gyri and sulci, in both cerebral hemispheres. Histogram plots show normalized mean ± SD (N=18).

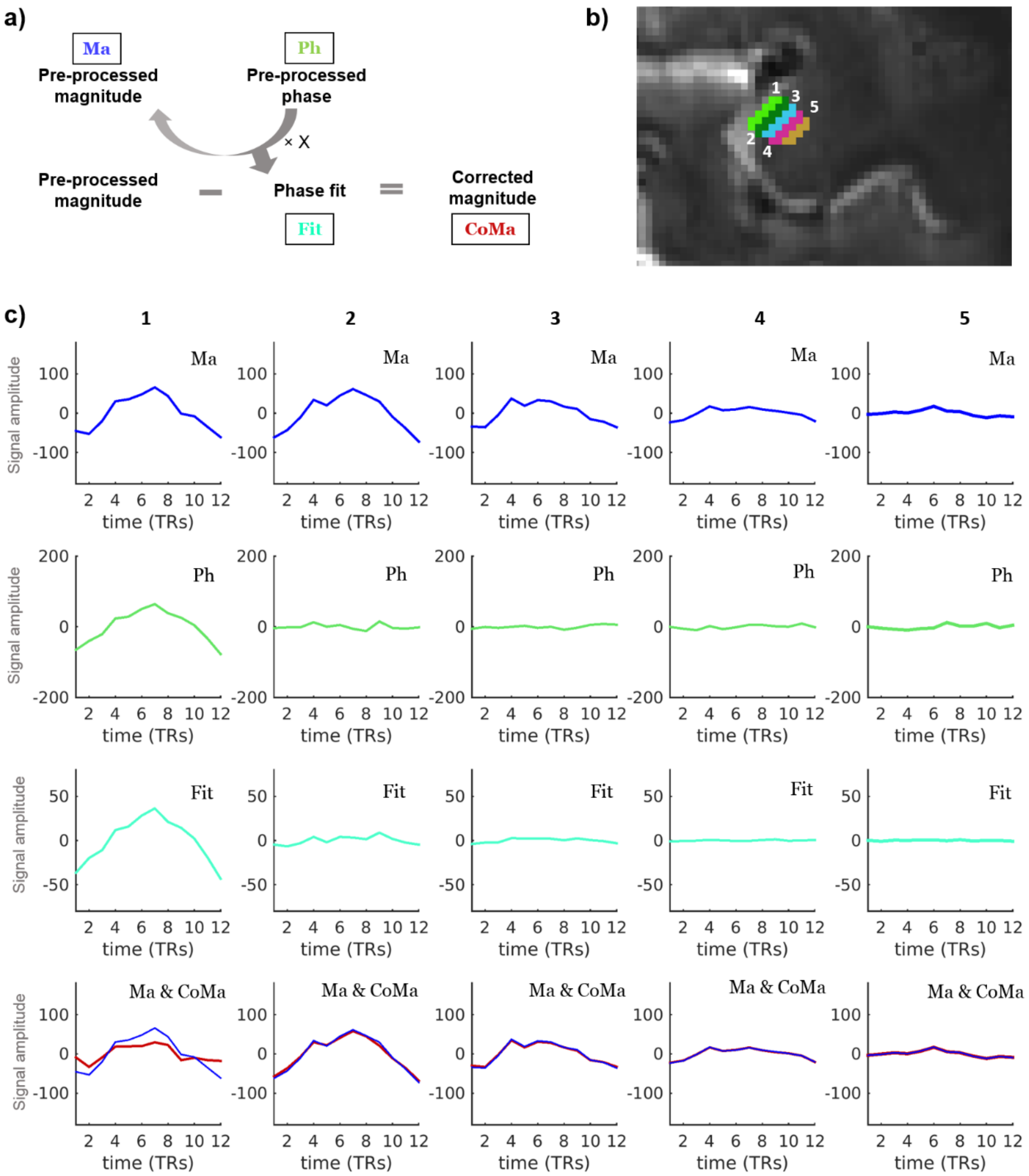


**Figure S3. Phase-based de-vein procedure. a)** Summarized procedure for the voxel-wise de-vein based on phase information. **b)** Location of regions selected to investigate the effect of the correction procedure, as depicted with different colors (1 to 5). “1” corresponds to voxels partially covering the CSF and “5” to deep gray matter/white matter. **c)** fMRI signal time courses, based on the average of 12 epochs of a motor task, for the five regions selected along the cortical depth and at different points in the correction procedure. The time-varying signal magnitude (top panels, in blue) was corrected by subtracting a weighted version of the phase (phase fit, in cyan). The bottom row shows the corrected time course (in red) as well as the original magnitude time course (blue line) for comparison. Note the relatively big effect of the procedure on the surface of the brain (bottom left panel in c)).

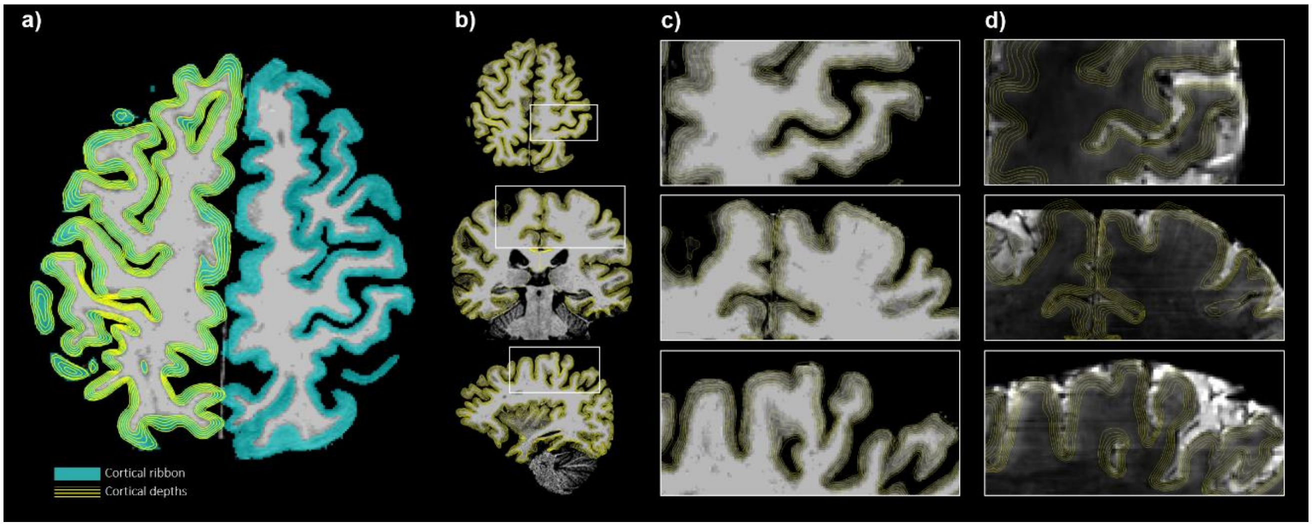


**Figure S4. Layer sampling. a)** An axial slice of the MP2RAGE showing the cortical ribbon segmentation (in blue) and the layers generated by Freesurfer in the left hemisphere (yellow lines). **b)** Example of axial, coronal, and sagittal MP2RAGE slices with the six cortical surfaces overlaid. **c)** Magnified view of the boxes in b). **d)** Layers overlaid on the mean functional image.

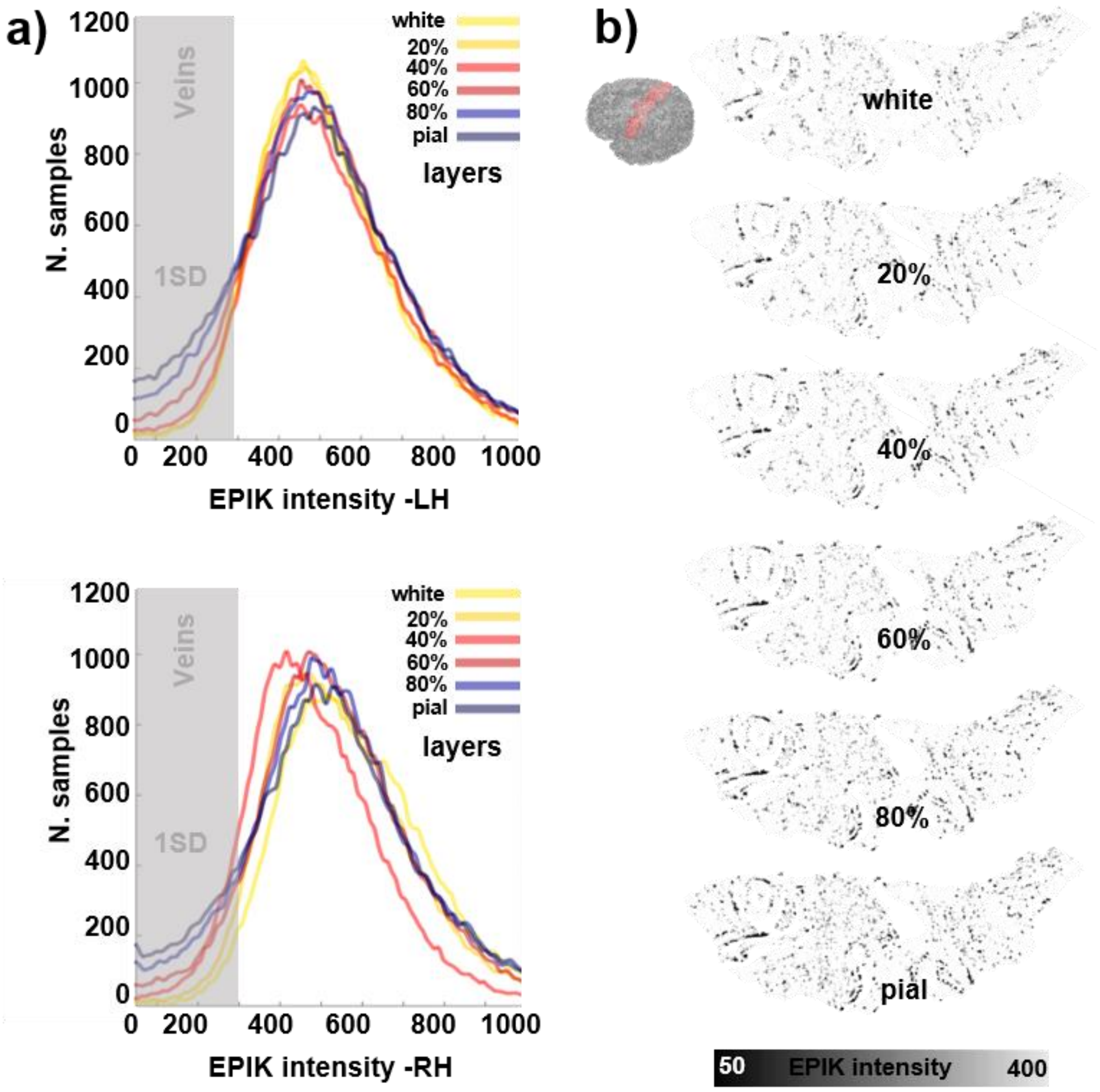


**Figure S5. Vein distribution across the cerebral cortex. a)** Histograms showing the average distribution of functional image intensity values across the vertices of surfaces generated at different cortical depths (N=18) for both cerebral hemispheres (LH/RH=left/right hemisphere). The gray-shaded area indicates the approximate range of values that were considered as veins in subsequent analysis (for each subject, a number of vertices equal to one standard deviation with the lowest intensity). **b)** An example of the venous locations across six different cortical depths in the left pre-central gyrus (flatten patches), identified as vertices with intensity below ~ 400 in the T2*-weighted image.

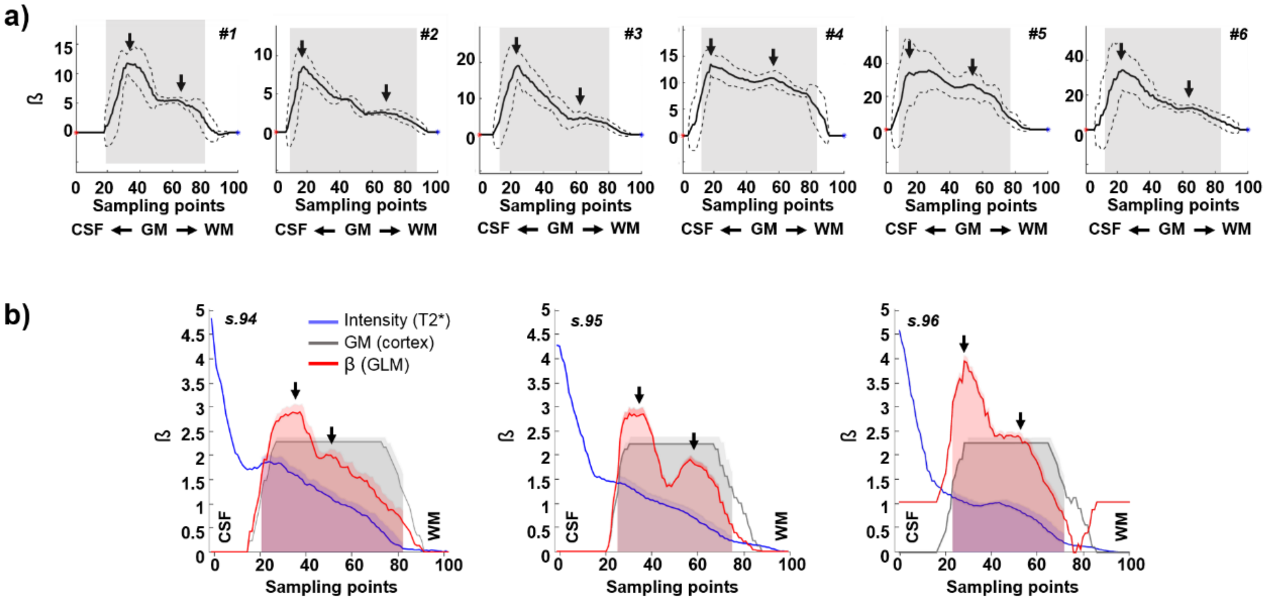


**Figure S6.** Examples of line activation profiles (computed as shown in Fig. 2) from **a)** six different subjects and **b)** three consecutive slices for one of the subjects. Sampling points 0 and 100 correspond to CSF and WM, respectively; sampling points along the shaded area correspond to cortical GM (cortical ribbon).

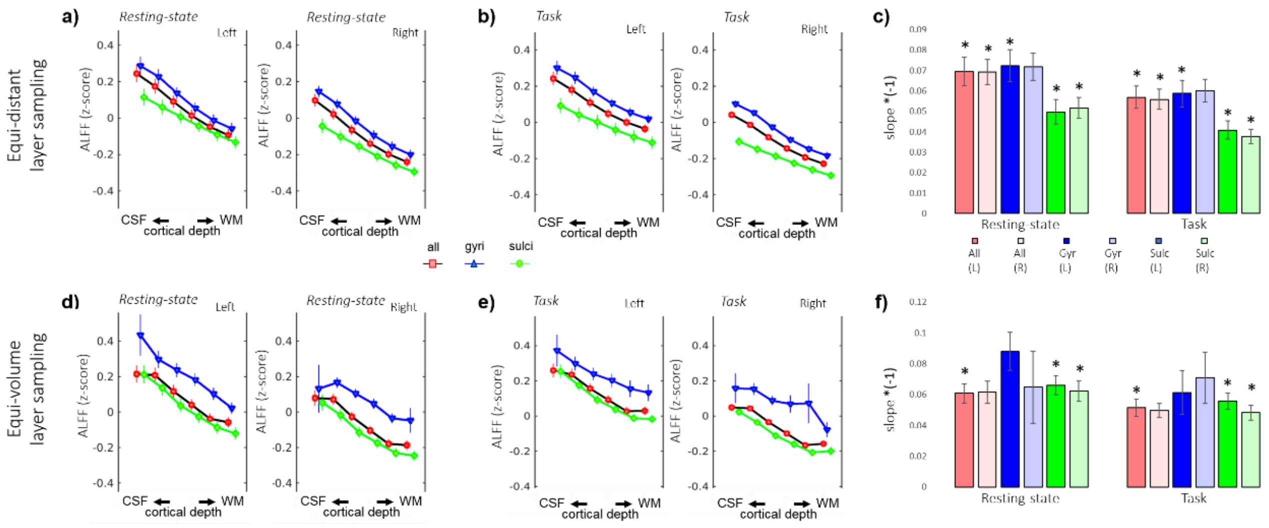


**Figure S7.** Assessment of ALFF along six cortical depths extracted following an equi-distance (**a-c**) or an equi-volume (**d-f**) model during resting-state and task and considering either all voxels (red), only gyri (blue), or only sulci (green). Panels **c)** and **f)** show the mean laminar slope for each condition (see legend below panel c)). Significant differences (p-value <0.05) encountered between resting-state, and task are marked with an asterisk (*) for each condition.

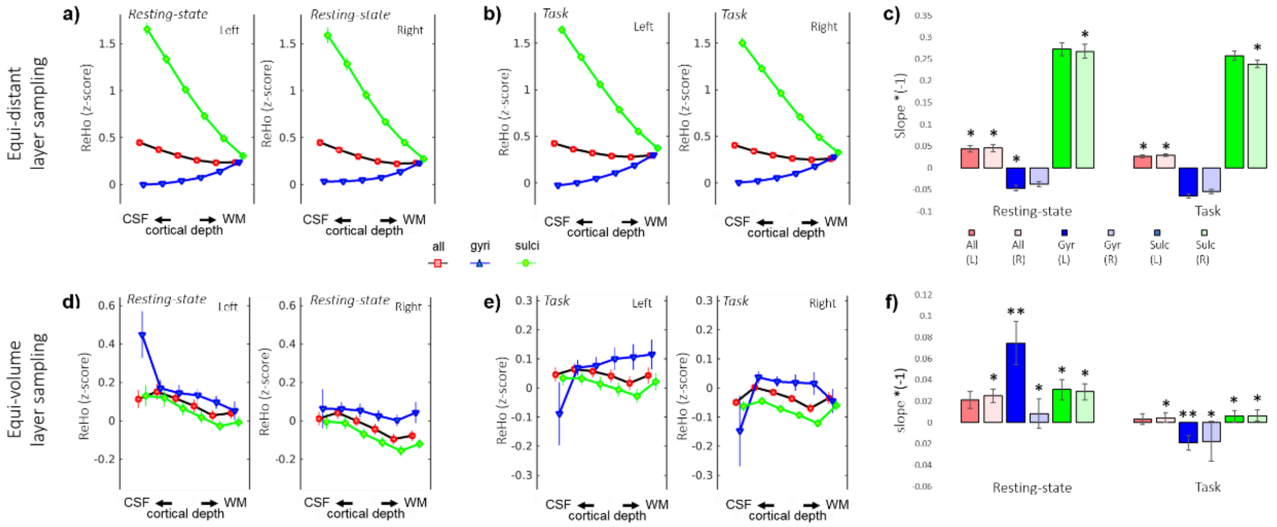


**Figure S8.** Assessment of ReHo along six cortical depths extracted following an equi-distance (**a-c**) or an equi-volume (**d-f**) model during resting-state and task and considering either all voxels (red), only gyri (blue), or only sulci (green). Panels **c)** and **f)** show the mean laminar slope for each condition (see legend below panel c)). Significant differences (p-value <0.05) encountered between resting-state and task are marked with an asterisk (*) for each condition.

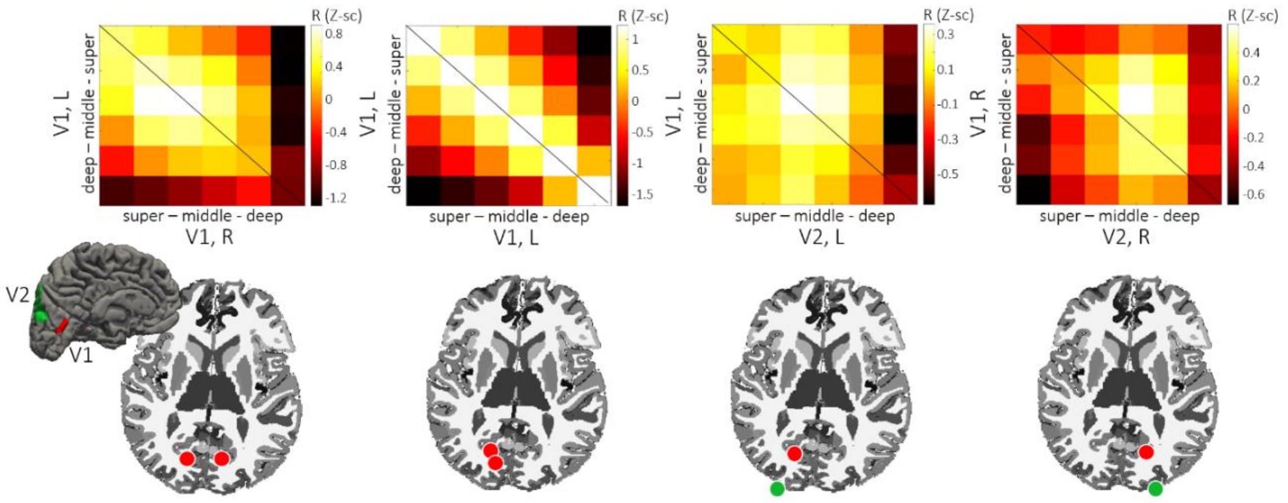


**Figure S9. Laminar connectivity in the visual system.** Connectivity evaluated as temporal correlation (R, z-scores) between six cortical depths of V1 and V2. V1: primary visual cortex; V2: secondary visual cortex.

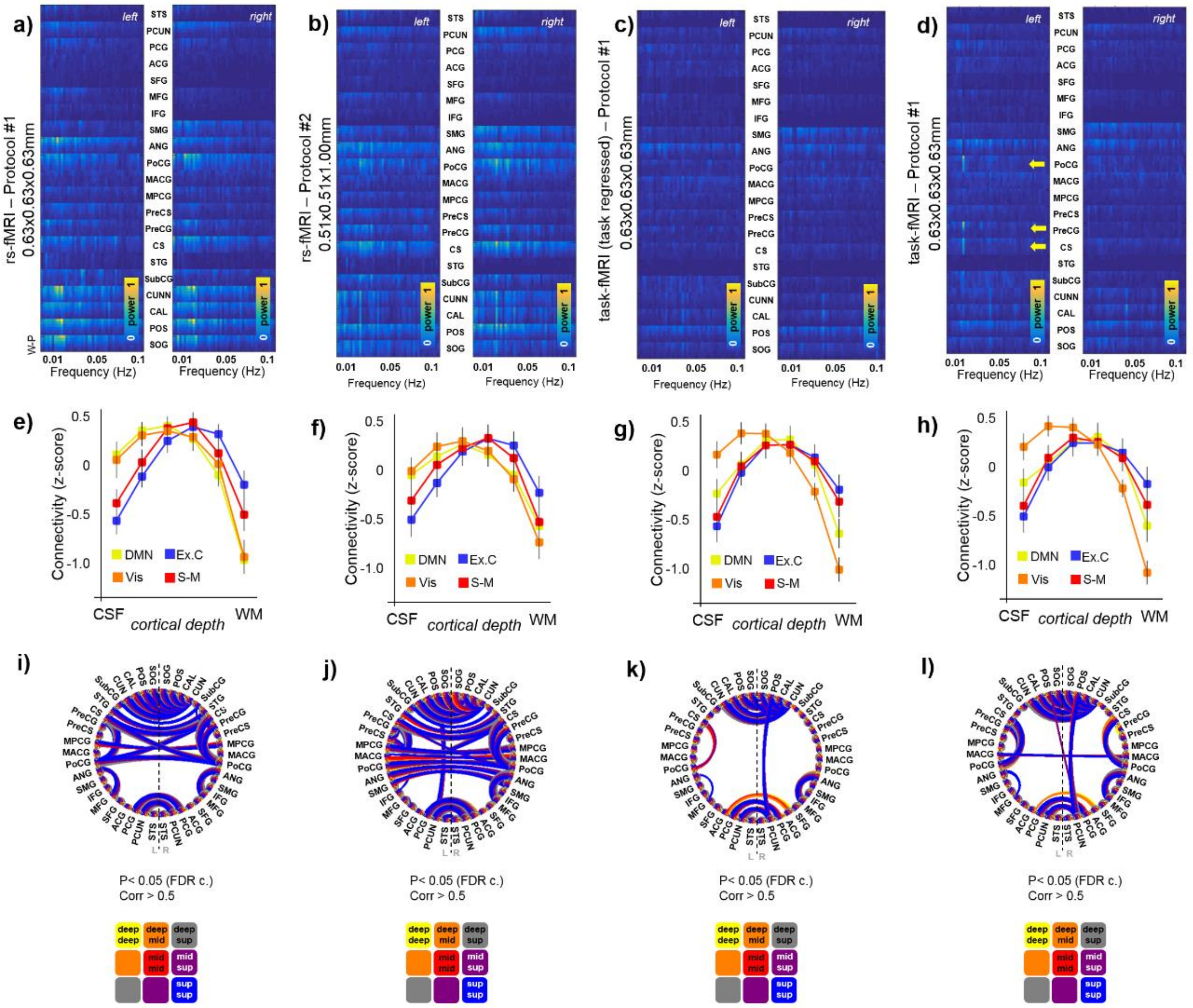


**Figure S10. Cross-scan data comparison.** For the three scans acquired from each subject, and in the case of the task fMRI, for data with and without regression of the task paradigm, the figure shows the following metrics: mean normalized frequency-power decomposition across six cortical depths in 21 ROIs (**a, b, c & d**), mean connectivity per network (color-coded) and cortical layer (x-axis) (**e, f, g & h**), and the whole-brain ROI/layer connectivity graph (**i, j, k & l**). N=13, 11, and 12 for protocol-1 resting-state, protocol-2 resting-state, and protocol-1 task fMRI with and without regression of the task paradigm, respectively.

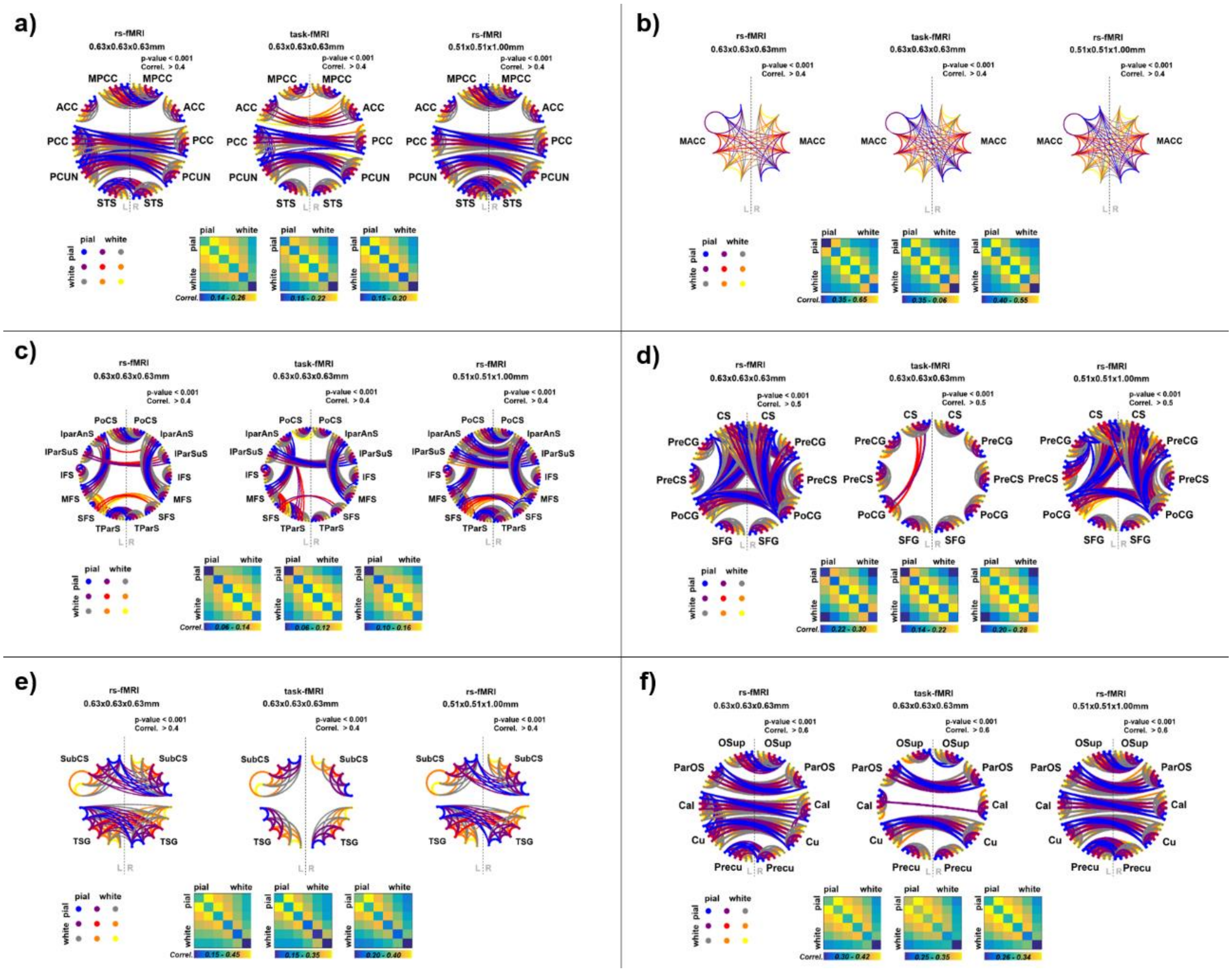


**Figure S11. Reliability of the resting-state networks.** The figure shows the group-connectivity graph and average layer-to-layer connectivity matrix for different resting-state networks: **a**) default mode, **b)** salience, here limited to the anterior mid-cingulate cortex due to incomplete coverage of the insula with protocol-1 in some volunteers), **c)** executive control, **d)** sensory-motor, **e)** auditory and **f)** visual, computed from the three different scans (from left to right: protocol-1: rest-, protocol-1: task (after regressing out the task paradigm), protocol-2: rest) across 13 subjects. The color of the lines indicates which pair of layers are correlated (see the color legend on the left below the graphs).

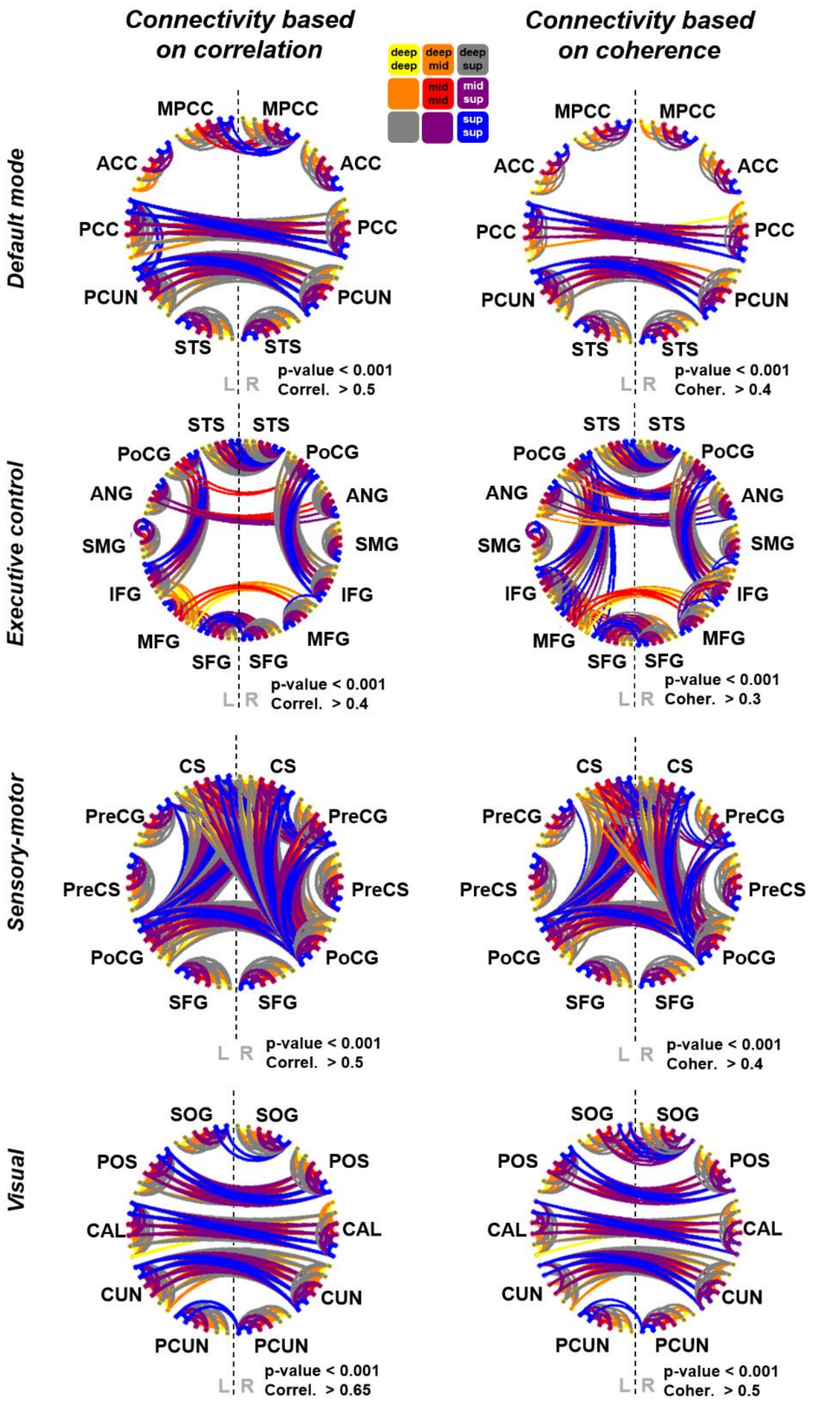


**Figure S12. Comparison between temporal-correlation-based and coherence-based connectivity results.** The four pairs of connectivity graphs show the most relevant connections based on a temporal correlation analysis (left) and a coherence analysis (right). Note the similarity between both sets of results.

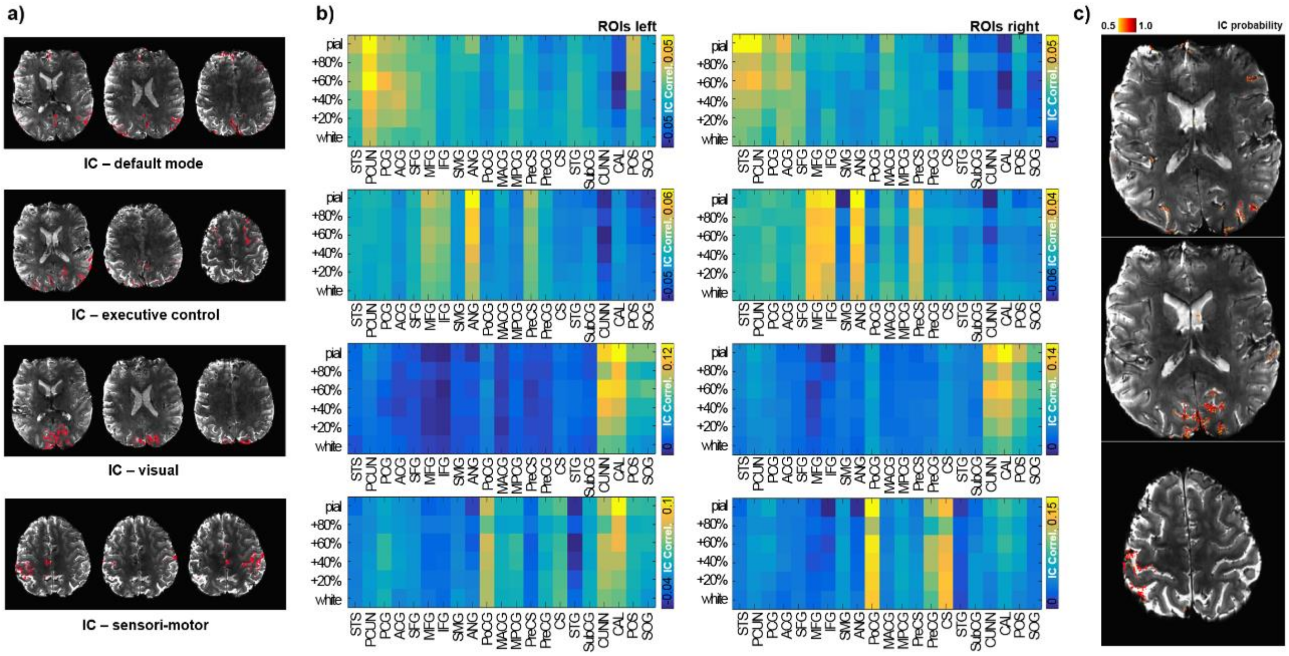


**Figure S13. Laminar detection of the volume-based independent components. a)** representative slices highlighting the voxels associated with four different independent components ("IC") corresponding to four functional networks. **b)** each color-coded panel shows the average temporal correlation between the time course represented by each independent component identified in volume-space and the time courses of vertices within 18 selected ROIs at six different cortical depths (mean correlation, N=13). Panels on the left and right sides represent the laminar correlation analysis for ROIs on the left and right hemispheres, respectively. Note the predominantly higher correlation values (i.e., higher contribution to a particular functional network) of superficial layers. **c)** Three examples of networks (executive control, visual, and sensory-motor) identified with ICA in a volunteer, localized in an axial slice, and magnified for visual inspection.

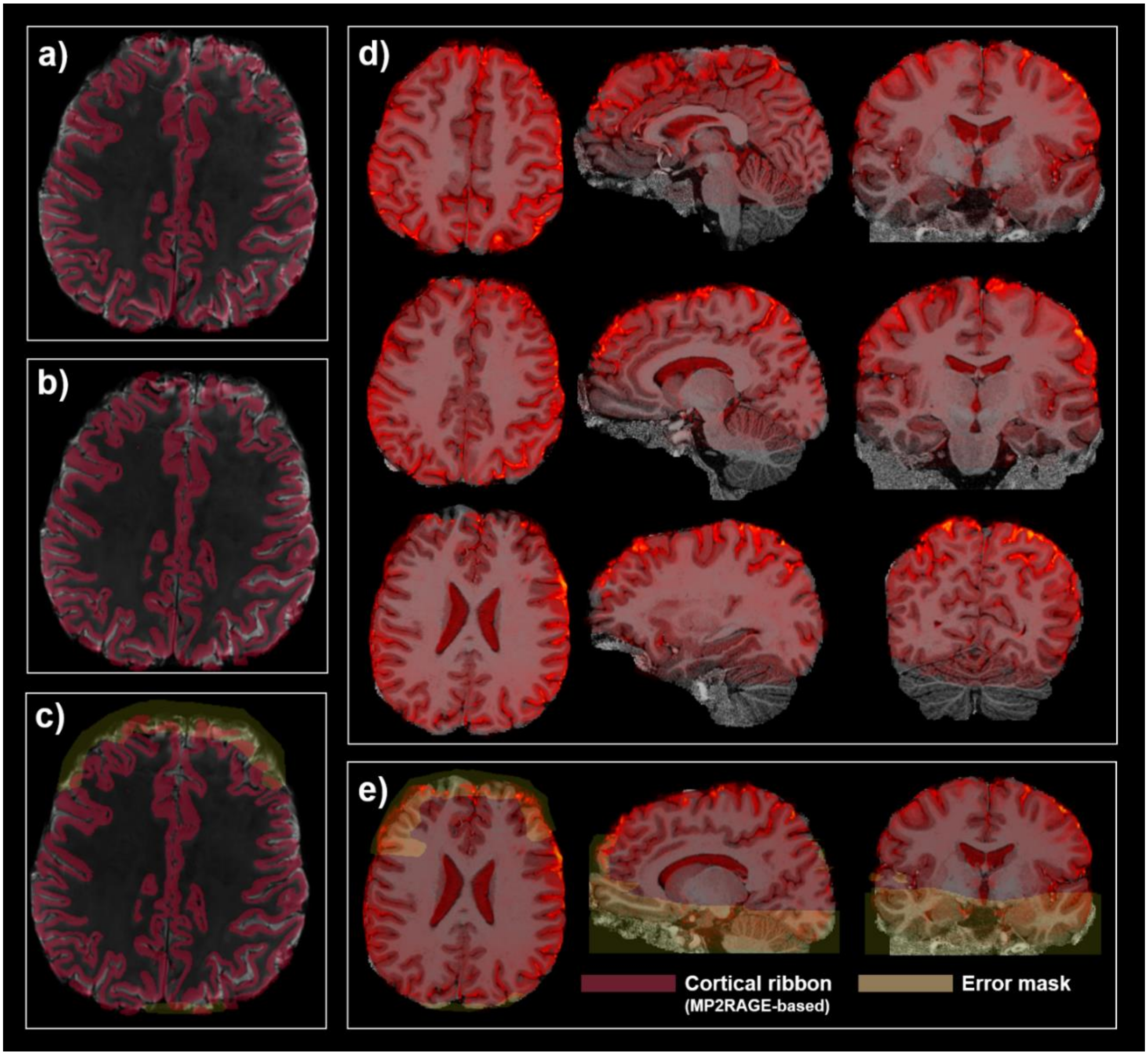


**Figure S14. TR-external EPIK-MP2RAGE co-registration. a)** Cortical ribbon (dark red) overlaid on a mean TR-external EPIK axial slice before co-registration. **b)** Co-registered cortical ribbon overlaid on the mean TR-external EPIK. **c)** Cortical ribbon overlaid on the co-registered TR-external EPIK scan and error mask (yellow) drawn over misaligned areas of the frontal lobe. **d)** Multiple slices showing the co-registration of the mean TR-external EPIK and MP2RAGE using MP2RAGE as the underlay and a semi-transparent heat-scale TR-external EPIK as the overlay. Note the near-perfect match between the dark CSF of the MP2RAGE and the bright CSF of TR-external EPIK. **e)** Example of co-registered slices and error-mask.

#### References used in the supplementary material
